# Dissecting the Determinants of Domain Insertion Tolerance and Allostery in Proteins

**DOI:** 10.1101/2023.04.11.536407

**Authors:** Jan Mathony, Sabine Aschenbrenner, Philipp Becker, Dominik Niopek

## Abstract

Domain insertion engineering is a promising approach to recombine the functions of evolutionarily unrelated proteins. Insertion of light-switchable receptor domains into a selected effector protein, for instance, can yield allosteric effectors with light-dependent activity. However, the parameters that determine domain insertion tolerance are poorly understood.

Here, we used an unbiased screen to systematically assess the domain insertion permissibility of several evolutionary unrelated proteins. Training machine learning models on the resulting data allowed us to dissect features informative for domain insertion tolerance and revealed sequence conservation statistics as the strongest indicators of suitable insertion sites. Finally, extending our experimental pipeline towards the identification of switchable hybrids resulted in opto-chemogenetic derivatives of the transcription factor AraC that function as single-protein Boolean logic gates. Our study reveals determinants of domain insertion tolerance and facilitates the engineering of switchable proteins with unique mechanistic properties.

## Introduction

The recombination of protein domains is an important driver of evolution. It allows nature to repeatedly build on the same set of stable protein folds and their corresponding functions, while enabling evolutionary innovation by exploring novel combinations and interdependencies thereof^1, 2^. This observation has inspired protein engineering approaches that combine evolutionary unrelated protein domains into single polypeptide chains, thereby creating hybrid proteins with new-to-nature properties^3–7^. From a synthetic biology perspective, a particularly interesting strategy is the insertion of receptor domains into effector proteins with the aim to allosterically couple the effector conformation to the receptor state^3, 5, 8^. Receptor activation, e.g. via chemicals or light, will induce an allosteric signal relaying to the effector’s active site (e.g. a catalytic surface or binding site), thereby enabling highly targeted control of the effector-mediated cellular function.

Although a number of hybrid proteins have been created by domain insertion engineering over the past years, their rational design remains challenging and screening of larger libraries and iterative optimization is commonly required to obtain functional hybrids^9–13^. Importantly, the identification of an insertion site at which the fusion of two protein domains results in their functional coupling and does not irreversibly interfere with the activity of either protein part represents a largely unsolved problem. These persisting challenges can be explained by our limited understanding of the structural and biophysical requirements and constraints that generally determine suitable domain insertion sites.

Advances in the generation of comprehensive domain insertion libraries via transposon-^12, 14^ or oligonucleotide pool-based cloning^15^, as well as the coupling of fluorescence-activated cell sorting (FACS) to deep sequencing, facilitate the efficient generation and subsequent investigation of larger domain insertion datasets^11, 12, 16^. Employing such experimental approaches, recent studies investigated the impact of domain insertion on the membrane localization of potassium ion channels^16, 17^. Using the resulting data to train random forest models, the authors analyzed biophysical properties that contribute to domain insertion permissibility in ion channels^17^. This previous research was centered around a single type of membrane proteins as well as the impact of domain insertion on subcellular protein localization. To render domain insertion engineering a broadly-applicable strategy, however, studying the domain insertion tolerance at the functional level as well as deciphering the determinants of functional coupling between re-combined protein domains will be essential.

Here, we set out to broaden the understanding of domain insertion requirements in diverse protein classes. Towards this goal, we inserted up to five structurally and functionally unrelated domains into several different, unrelated candidate effector proteins covering nearly all possible sequence positions. Using gene circuits that relay effector activity to a fluorescent readout, the resulting, comprehensive libraries of protein hybrids were screened for active variants by FACS and subsequent next-generation sequencing (NGS). Training of machine learning models on the resulting datasets allowed us to dissect parameters that affect domain insertion tolerance and revealed sequence conservation statistics as the most powerful predictors for domain insertion success. Finally, extending our experimental pipeline towards the screening of engineered, switchable effector variants yielded two potent optogenetic derivatives of the *E. coli* transcription factor AraC that function as single-protein chemo-optogenetic Boolean logic gates.

## Results

### A functional FACS-NGS screen of domain insertion tolerance

To elucidate the domain insertion tolerance within an evolutionarily and functionally diverse set of effector proteins, we first constructed comprehensive insertion libraries. The libraries comprised of effector proteins carrying insert domains at all possible sequence positions (Fig. 1a). Four structurally unrelated proteins that are widely applied in synthetic and cell biology were chosen as effector protein scaffolds: the transcription factor AraC, the recombinase Flp, a previously described variant of the TVMV protease^18^ and ơ-factor F (SigF) from *Bacillus subtilis* (Fig. 1b). Protein hybrid libraries were generated via saturated programmable insertion engineering (SPINE) for all four candidates using the PDZ domain from murine α1-syntrophin as insert (Fig. 1b)^15^. With its small size of 86 amino acids, its globular fold and the N- and C-terminus located in close proximity, the PDZ domain is ideally suited for domain insertion screening^11^. Further, to elucidate how the domain identity would affect the functionality of the resulting protein hybrids, four additional insert domains of varying size and structure were selected and fused at all possible sequence positions into one of the candidate proteins, AraC. These included the *As*LOV2 (*Avena sativa*) domain, the estradiol binding domain from human estrogen receptor-α (ERD), an enhanced yellow fluorescent protein (eYFP)^19^ and the synthetic rapamycin receptor uniRapR^20^. Following the construction of all eight libraries, a near complete coverage of all possible insertion sites was observed by deep sequencing (Supplementary Fig. 1).

**Fig. 1.**
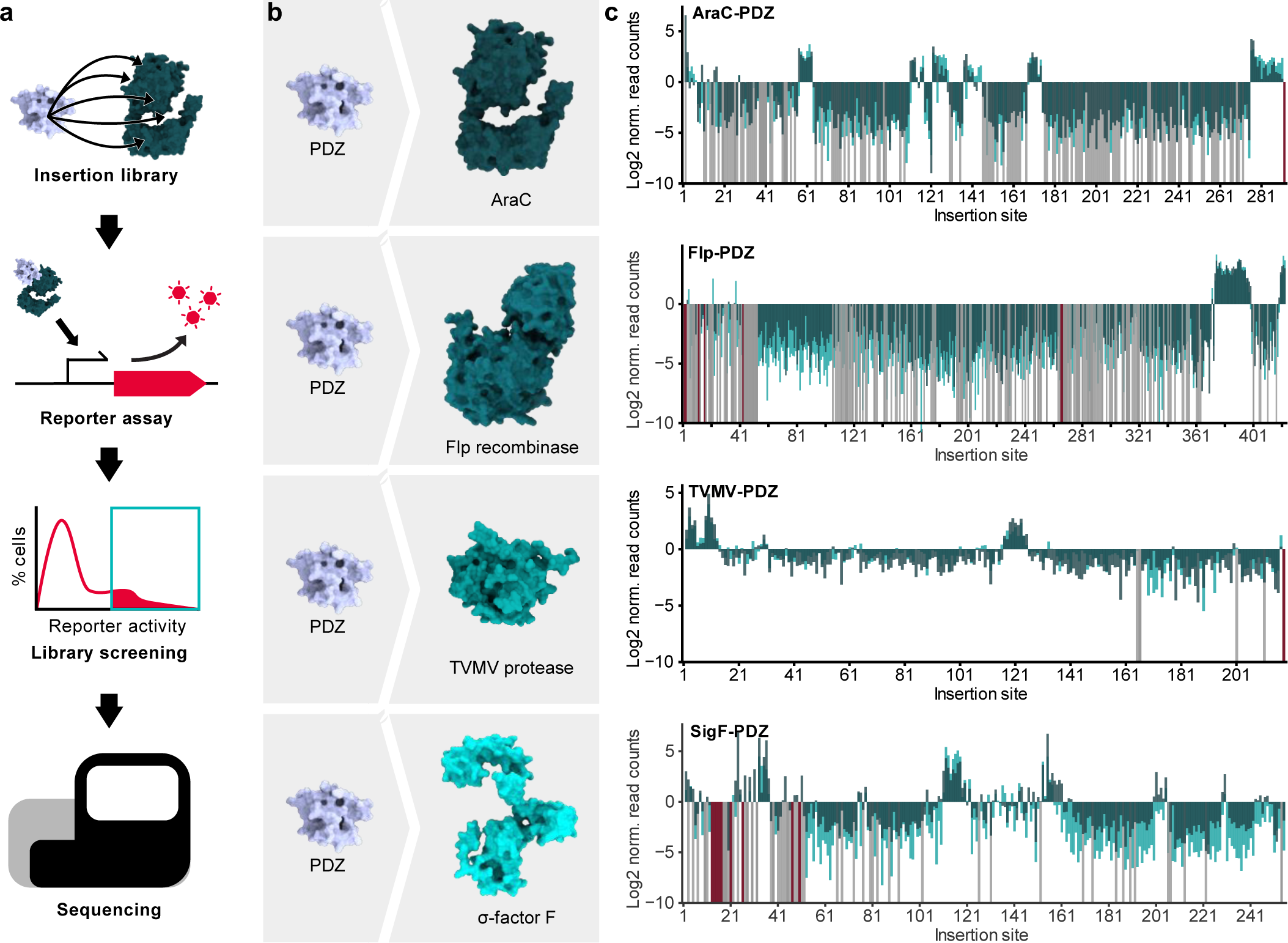
Domain insertion profiling of functionally and structurally diverse proteins. **a**, Flow chart of the domain insertion screening workflow. **b**, Overview of the screened PDZ-domain insertion libraries. The depicted structures of the parent proteins are AF2 predictions. PDB-ID of PDZ: 1Z86. **c**, Enrichment score histograms for the different candidate proteins are shown. The Log2 norm. read counts correspond to the fraction of reads after enrichment normalized to the fraction of read counts within the initial library. Data from the four candidate proteins AraC, Flp, TVMV protease and SigF with PDZ domain inserts are shown. Enrichments are mapped to the respective insertion site as indicated by the position of the acceptor proteins preceding the insertion. Light green, dark green: individual replicates. Grey: variants with zero reads after enrichment. Red: variants missing in the initial library. Insertion sites correspond to residues preceding the inserted domain.

To enable functional screening of these libraries in *Escherichia coli*, we next created reporter gene circuits that robustly couple the activity of the effector protein to the expression or stability of a red fluorescent protein (RFP) (Extended Data Fig. 1, Methods). We then co-transformed *E. coli* Top10 cells with the reporters and their corresponding effector-insert hybrid libraries, followed by analysis of the reporter activity via FACS. Fluorescence histograms of the initial libraries showed a large fraction of non-functional hybrid protein candidates as indicated by a large proportion of non-or low fluorescent cells (Extended Data Fig. 1). Still, a small but considerable fraction corresponding to medium to high fluorescent cells and hence active protein hybrids was observed. Sorting this fraction resulted in a clear enrichment of cells expressing high RFP levels in case of AraC and SigF and less pronounced, but still visible enrichments in fluorescent cells for Flp and the TVMV protease (Extended Data Fig. 1). Quantitative differences between the four effector library pools were caused by varying proportions of active versus inactive hybrid protein candidates in the initial libraries as well as differences in the dynamic range of the reporter assays (Extended Data Fig. 1, controls). To ensure a significant enrichment of active variants, we sorted each library in two consecutive rounds. Next, we assessed enrichment or depletion of each individual domain insertion variant in the sorted libraries by adapting the previously published DIP-seq pipeline^12^. In short, the fraction of read counts corresponding to a variant after enrichment were normalized by the fraction of read counts from the initial library and the resulting scores were log2-scaled. Variants that went extinct during sorting were automatically assigned the value −10, which was in the range of the strongest measured depletions. To ensure reproducibility of the workflow, the whole screening and sequencing process was performed in two independent replicates.

Results from different replicates correlated well, with Pearson’s r > 0.8 in all cases except one (Pearson’s r for TVMV-PDZ = 0.65), while the level of enrichment/depletion differed between replicates for individual variants (Supplementary Fig. 2). As cross-validation of our enrichment and analysis pipeline, we experimentally measured the activity (RFP expression) for a set of hybrids individually and compared it to the variant enrichment scores obtained by NGS. As expected, a drastic difference in activity between the enriched and the depleted variants was measured in most cases (Supplementary Fig. 3). For the following analysis the mean of the two biological replicates was used.

### Domain insertion permissibility is sequentially and structurally clustered

Mapping the enrichment scores of the PDZ insertion libraries to the amino acid sequences of the respective, four effector proteins revealed that positions tolerating insertions occurred in clusters spanning regions of ∼10-30 consecutive amino acids (Fig. 1c). Insertion tolerance thus appears to be regionally confined, rather than being determined by features of individual residues or positions. Roughly 80 % of the insertions within each protein were depleted, i.e. they do not tolerate domain fusion (Fig. 1c). Moreover, the number of clusters with enrichments differed substantially between the insert domains tested for the AraC effector (Extended Data Fig. 2). For the LOV2 insert domain, we observed several insertion-permissive regions throughout the sequence of AraC comparable to those for the PDZ insert. In contrast, the other three insert domains were enriched at substantially fewer positions, mainly at the C-terminus of AraC. As LOV2 and PDZ are considerably smaller (<150 AA) than the other tested domains, insert size appears to be a determining factor for insertion tolerance. Interestingly, we hardly observed insertion sites selective for just one specific insert domain. This indicates that domain insertion permissibility is a general property of protein regions rather than a lock-key relation between an insertion site and an individual insert domain.

Next, we mapped the enrichment scores onto structures of the respective effector proteins. To this end, we used Alphafold2 (AF2)-predicted protein structures, as well as experimentally resolved full length structures if available^21, 22^ (Fig. 2a-d, Extended Data Fig. 3 and 4, Supplementary Fig. 4). Importantly, the predicted structures were generally in excellent agreement with the available experimentally validated (partial) folds (Supplementary Fig. 5). Structural analysis revealed strong depletions around functionally critical regions, such as the DNA- and arabinose-binding sites of AraC, the catalytic center of the Flp recombinase or the DNA-binding region of SigF (Fig 2a-d, Extended Fig. 3, Supplementary Fig. 4). For TVMV protease, depletions within the hydrophobic core and around the active site were observed, albeit trends were overall less pronounced for this candidate protein (Fig 2c, Extended Fig. 3c). Interestingly and in contrast to common assumptions underlying domain insertion engineering strategies, no clear enrichment at surface-exposed unstructured loops could be identified for any of the candidates. Rather, insert sites were observed at similar frequency in helices, sheets and loops (Fig 2a-d).

**Fig. 2.**
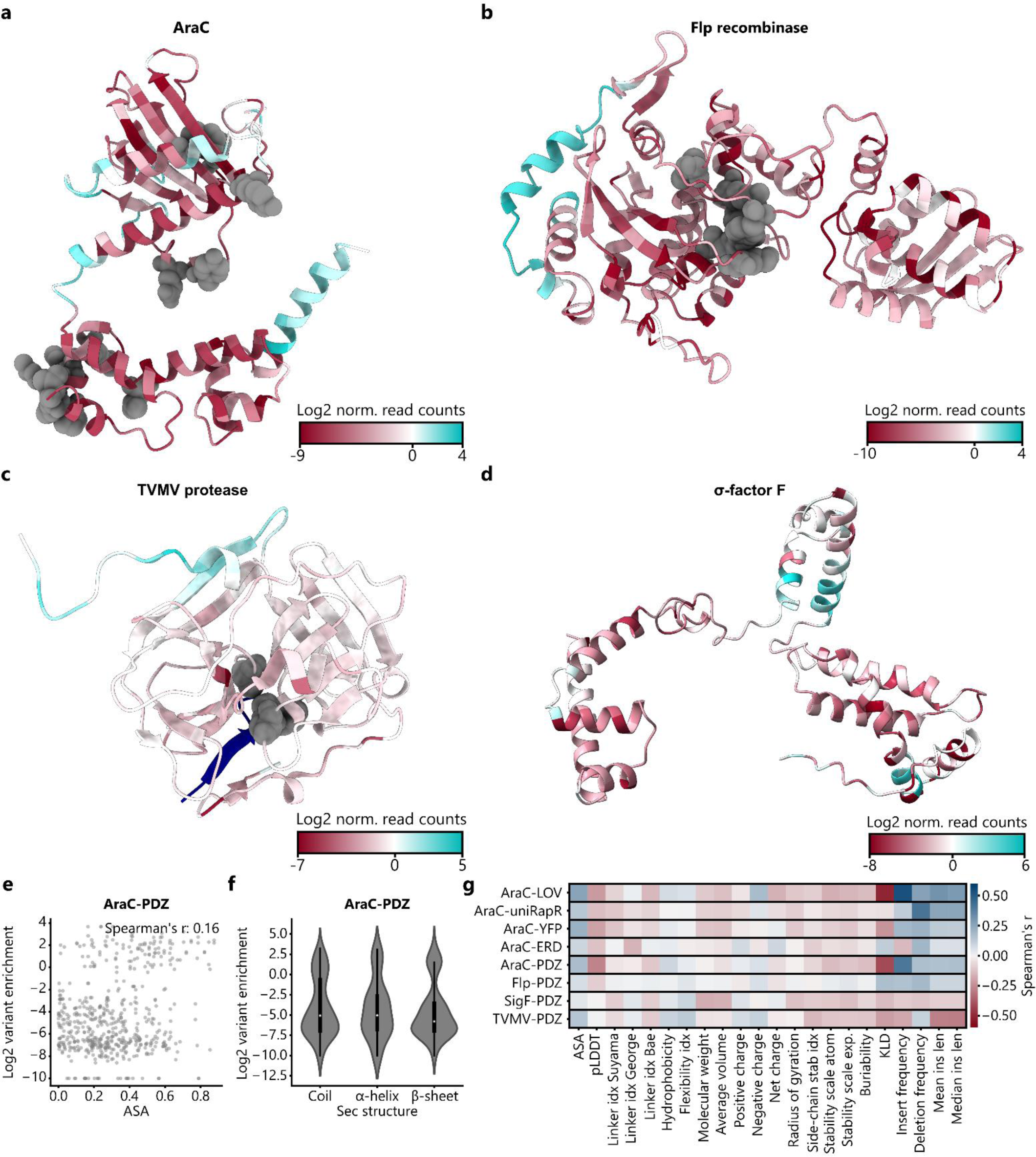
Secondary structure and amino acid features alone do not explain the experimentally observed domain insertion patterns. **a,** Domain insertion permissive positions are clustered at diverse, locally confined surface sites. The insertion scores from the PDZ libraries are mapped onto the AF2 structure predictions of the candidate proteins namely AraC (**a**) and Flp recombinase (**b**), the crystal structure of the TVMV protease (PDB-ID: 3MMG) (**c**), and an AF2 structure prediction of SigF (**d**). Functionally critical residues of AraC, Flp and the TVMV protease are indicated in grey. **e**, Correlation between variant enrichment and the average surface exposed area (ASA) of the residues neighboring an insertion site are plotted for AraC-PDZ. Spearman’s r is indicated. **f,** Violin plot of the insertion score distribution with respect to different secondary structure elements is shown for the AraC-PDZ insertion library. For each insertion site, the secondary structure assignment of the amino acids prior and after the insertion were considered. The IQR is marked by the box and the median is represented by a white dot. Whiskers extend to the 1.5-fold IQR or to the value of the smallest or largest enrichment, respectively. **g**, Spearman correlations between all datasets and diverse positional features are shown (Extended Data Table 1. Linker idx: Different amino acid specific linker propensity indices that were reported by the indicated authors.

Next, to quantitatively analyze these qualitative observations, we correlated the measured enrichments with a set of basic positional properties such as the average solvent accessible area (ASA), secondary structure and amino acid identity of the residues neighboring a respective insertion site (Fig. 2e, f, Supplementary Fig. 6 and 7). Of note, none of these basic properties explained the observed enrichments. In order to obtain a more comprehensive overview of protein features that could affect domain insertion success, a larger set of position-specific features was gathered (Extended Data Table 1, Methods). Further, these comprised a number of biophysical amino acid properties, fetched from the “AAindex” database^23, 24^, as well as several previously published linker propensity indices^25–27^. These indices describe to which extend amino acids tend to be present in inter-domain linkers. Regions with high linker propensities are commonly expected to be well suited for the insertion of domains. Further, we included the pLDDT confidence score from AF2 models, which was previously shown to correlate with intrinsically disordered sites^28^. Moreover, the Kullback-Leibler divergence (KLD), a measure for sequence conservation, was extracted from multiple sequence alignments of the candidate protein with natural homologues. Finally, additional scores, such as the frequency of insertions and deletions at every position in evolutionary related sequences, were included (refer to Methods). Spearman correlations between all enrichment scores for the screened libraries and each feature revealed overall weak trends, with the majority of the correlation coefficients lying in the range between −0.2 and 0.2 (Fig. 2g, Supplementary Fig. 8). This observation is in agreement with previous results in the context of ion channels^16, 17^. Additionally, we confirmed that AF2-based structure predictions of insertion variants could not explain the observed enrichment trends (Supplementary Note 1, Supplementary Fig. 9 and 10).

### Machine learning reveals statistical features predicting domain insertion tolerance

The absence of any clear correlation between the experimental data and positional protein properties raised the question if a combination of the above features would enable the prediction of domain insertion tolerance. To address this question, machine learning models were trained on the entirety of the gathered insertion site properties in combination with amino acid identity and secondary structure information as additional features. The learning objective was to discriminate between enriched sites that tolerated the insertion of a domain versus depleted positions, as these states appeared to be well separated in the data (Fig. 1c). We trained gradient boosting classifiers^29^ for each protein using five-fold cross-validation. The models reached surprisingly good performances on datasets derived from individual candidate proteins ranging from a mean area under the receiving operator characteristic (AUROC) of 0.72 for SigF-PDZ to 0.92 for Flp-PDZ (Fig. 3a, Extended Data Fig. 5). The corresponding average precisions (AP) ranged from 0.41 (SigF-PDZ) to 0.82 (AraC-PDZ) (Fig. 3a, Extended Data Fig. 5). The lower AP values are caused by the high proportion of negative labels in the respective datasets. Encouraged by these results, we optimized the model on a complete training set including all four proteins, which resulted in a mean AUROC of 0.84 and a mean AP of 0.54 (Fig. 3b). To place the classifiers performance into context, we compared it to several benchmarks on a previously withheld test set. These included a random choice baseline, and the use of individual features as predictors. Our classifier exhibited highly improved predictive power as compared to all individual features, reaching an AUROC of 0.85 and an AP of 0.56, suggesting that the entirety of input features implicitly provided the information necessary for successful prediction of domain insertion tolerance (Fig. 3c, Supplementary Fig. 11).

**Fig. 3.**
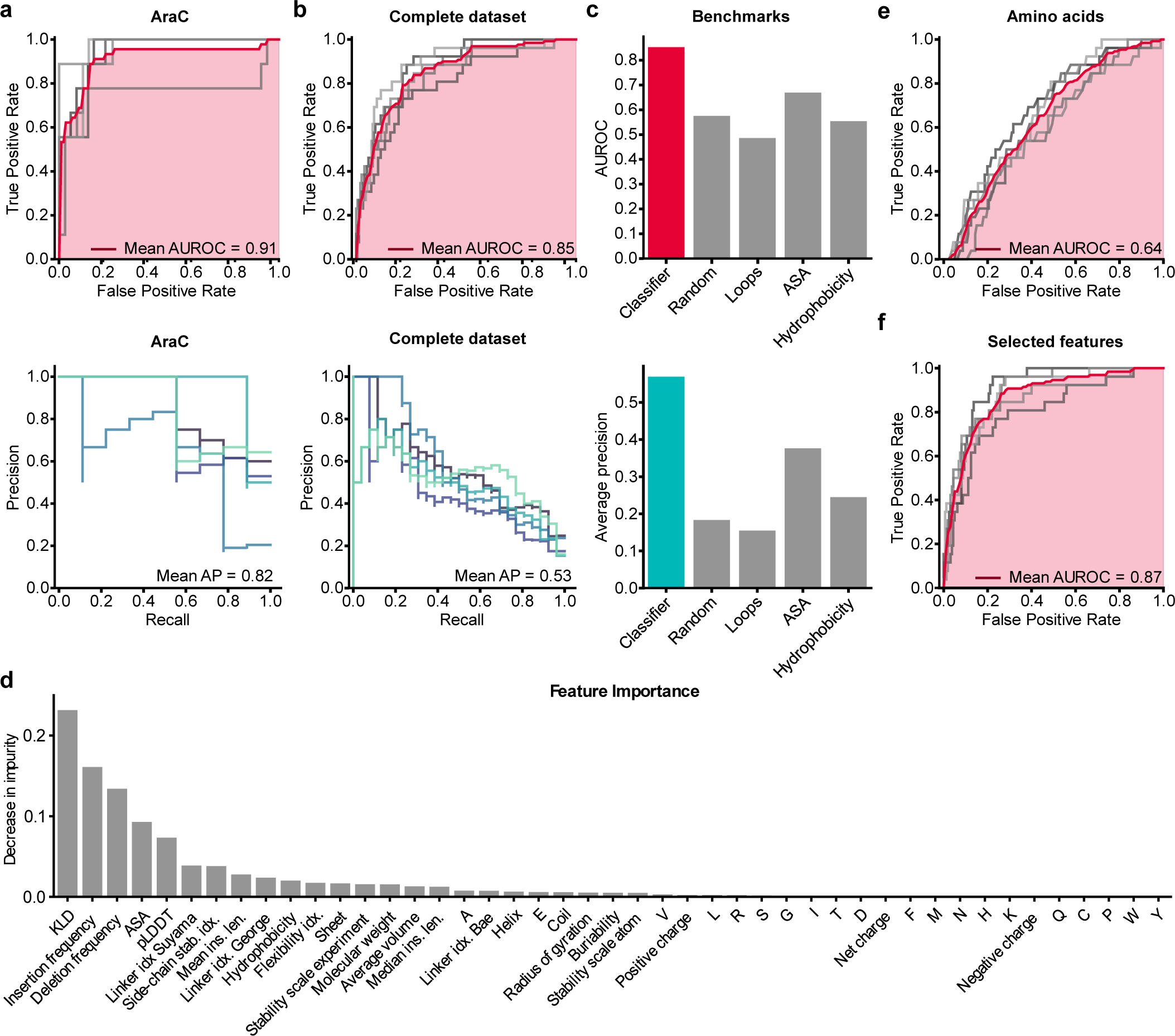
Gradient boosting classifier models reveal parameters informative of domain insertion tolerance. **a**, **b** ROC of the model, trained on the AraC-PDZ dataset (**a**) or the combined PDZ datasets of all candidate proteins (**b**) with five-fold cross-validation (Top). Precision-recall metrics for individual folds are shown (Bottom). The mean average precision is indicated. **c**, The AUROC and average precision of the trained classifier and different benchmarks are shown. The values were calculated from a previously withheld test set. **d**, Bar plot indicating the Gini importance of each feature for the model trained on the full dataset. **e**, The ROC metric of a gradient boosting model that was trained exclusively on the amino acid identities is shown. **f**, ROC of a model that was trained on a subset of features comprised of Deletion frequency, KLD, insert frequency, mean insertion length, the linker propensity index by Suyama^25^ and the pLDDT score from AF2 structure predictions. **b**, **e**, **f**, The ROC is depicted for individual folds in grey and the mean ROC in red. The mean AUC is marked in red. Precise values are indicated.

Finally, we aimed at identifying the key features most informative for the prediction of domain insertion tolerance. To this end, the importance of individual features for model performance was assessed by measuring the permutation importance of each feature as well as its Gini importance^30^ (Fig. 3d, Supplementary Fig. 12a). Both measures indicated that most parameters were dispensable, while the alignment-derived properties were most critical for successful prediction. In that line, a model trained solely on information about the identity of insertion-adjacent amino acids did reach an AUROC of 0.64 (Fig. 3e). As a consequence, we depleted features from the input data in a stepwise manner, while ensuring the performance of the model did not decrease upon feature removal. Following this procedure, we were able to train a reduced model, only based on six features: KLD, deletion frequency, insertion frequency, mean insertion length, pLDDT and the linker index by Suyama et al.^25^. With an AUROC of 0.87 and an AP of 0.55, the reduced model performed as good as the original one trained on all features (Fig. 3f). Lastly, the feature importance analysis was repeated with the reduced model. Akin to the previous observations, KLD, insertion frequency and deletion frequency, i.e. evolutionary and statistical features derived from MSAs, were detected as most important parameters explaining domain insertion tolerance (Supplementary Fig. 12 b, c).

### Identification of potent light-switchable AraC variants

Up to this point, we focused on features determining the preservation of function upon domain fusion into an effector protein. Taking our experimental screening approach one step further, we next investigated to which extend insertions can mediate allosteric behavior, i.e. a functional link between an insert and the effector. Such switchable hybrids are of great interest for various applications in biology and bioengineering. Towards this goal, we re-visited our initial AraC-LOV hybrid library. The *As*LOV2 domain is known to reversibly unfold its two terminal helices in response to blue light (∼450 nm), a property that has been harnessed for the development of light-switchable effector proteins in optogenetics^3, 31^. It was hence interesting to explore, whether screening our comprehensive AraC-LOV library could readily reveal potent, optogenetic AraC variants.

We therefore repeated the screen for the AraC-LOV library, this time incubating the cultures under blue-light exposure prior to FACS sorting. The resulting variant enrichment was then compared to that of the same library sorted upon incubation of cultures in the absence of light (Extended Data Fig. 2). Globally, we observed a high similarity between the resulting enrichment scores for each position under both conditions (Fig. 4a, Extended Data Fig. 6a). However, a subset of regions showed significant differences between the enrichment scores obtained for the libraries cultured in the dark and light (Extended Data Fig. 6b-c). Strikingly, further analyzing the insertion variants in these regions revealed a plethora of presumably light-activatable as well as light-inhibited AraC-LOV hybrids corresponding to multiple different AraC insertion sites (Extended Data Fig. 6b-c).

**Fig. 4.**
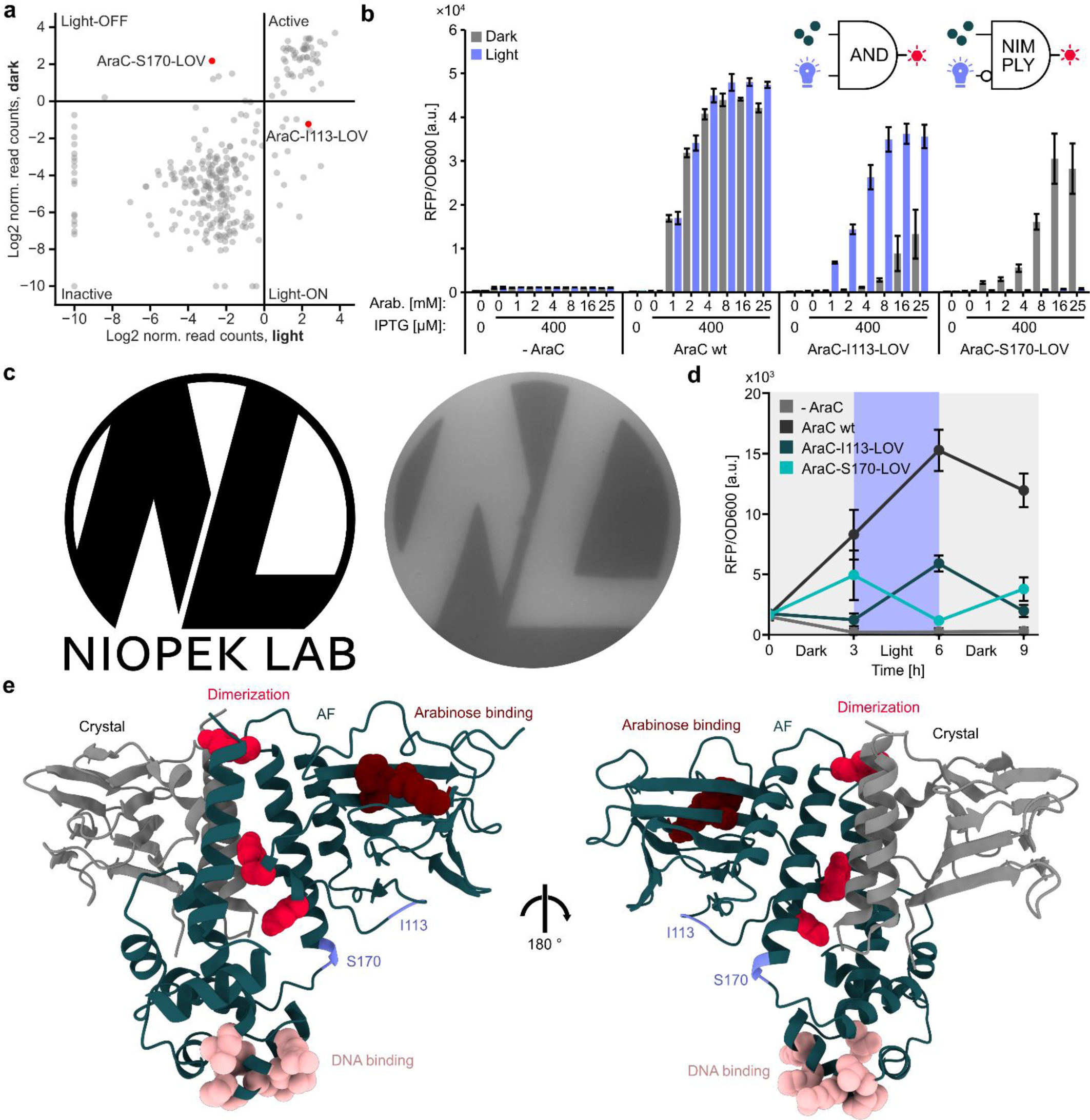
LOV2 domain insertion screening yields chemo-optogenetic AND and NIMPLY gates. **a**, Scatterplot showing the relation between the enrichment scores for the libraries incubated in the light and dark for each insertion variant. The two candidates selected for further characterization are marked in red and their names are indicated. **b**, Characterization of light-responsive AraC variants. Inducers were supplied in the indicated concentrations. The samples were incubated for 16 h under light exposure or in darkness, followed by plate reader measurements of reporter fluorescence (RFP) and OD600. Bars represent means from three independent replicates. Error bars show the SD. AraC-I113-LOV2 acts as a single-protein AND gate and AraC-S170-LOV2 as a NIMPLY gate. **c**, Agar photograph generated via an AraC-S170-LOV2 controlled RFP reporter. Top agar mixed with inducers and bacteria carrying an RFP reporter plasmid and the AraC-S170-LOV2 variants was plated on an ager plate, which also contained arabinose and IPTG. The plate was incubated overnight, while being illuminated through a photo-mask of the logo on the left (without the text). **d**, Cultures were inoculated from precultures carrying plasmids encoding an RFP reporter and the respective AraC variant into media carrying 400 μM IPTG and 25 mM arabinose. The cultures were incubated either in darkness or under blue-light exposure. At the beginning of the experiment and every three hours from then, RFP fluorescence and OD600 were measured, followed by 1:30 dilution in fresh media. Points represent the mean of n=3 biological replicates. Error bars indicate the SD. **e**, An AF2 prediction of the full-length AraC (green) is shown alongside the crystal structure (grey) of the arabinose binding domain. Relative positioning of the structures was obtained by superimposing the AF2 model onto a dimer crystal structure. Residues that bind to the operator are highlighted in pink, residues that are key for dimerization in presence of arabinose are shown in red, and the amino acids that are important for arabinose-binding in vermilion. The insertion sites corresponding to the AraC-I113-LOV and AraC-S170-LOV hybrids are marked in blue. PDB-ID: 2ARA.

From this set of optogenetic variants, we chose two AraC-LOV hybrids for further characterization, one light-ON switch carrying the LOV2 insertion behind I113 (AraC-I113-LOV) and a light-OFF switch with the LOV2 insertion behind S170 (AraC-S170-LOV). We then assessed the performance of these AraC-LOV hybrids using the previously established RFP transcription reporter in *E. coli* under varying arabinose concentrations, as well as light conditions. Interestingly, the activity of both AraC-LOV hybrids was co-dependent on the arabinose concentration and the light stimulus (Fig. 4b, Extended Data Fig. 6d). The AraC-I113-LOV samples showed a 23-fold increase in reporter expression upon illumination at an arabinose concentration of 4 mM. At higher arabinose concentrations, increasing fluorescence levels were also observed for samples incubated in the dark, indicating that the chemical inducer could, to some extent, override the light-mediated regulation. Vice versa, the AraC-S170-LOV samples showed efficient, light-dependent repression of reporter activity practically to baseline with a 43-fold switch in reporter activity at 16 mM arabinose. Moreover, the light-regulation was in this case not affected by high arabinose concentrations. The observed behavior establishes the AraC-I113-LOV and AraC-S170-LOV hybrids as single-protein Boolean logic devices capable of integrating light and arabinose as inputs and functioning as AND and NIMPLY gates, respectively (Fig. 4c-d, Supplementary Note 2).

Next, we investigated if these new optogenetic AraC variants facilitate spatiotemporal control of gene expression. Growing the AraC-S170-LOV reporter strain on agar while illuminating it through a photo-mask confined reporter RFP expression to light-shielded regions and hence resulted in display of the photomask shape on the fluorescent cell layer (Fig. 4c). Moreover, incubating AraC-S170-LOV and AraC-I113-LOV reporter strain cultures while alternating between light and dark conditions resulted in reporter expression oscillation, the phase of which depended on the AraC-LOV variant used (Fig. 4d). Taken together, the results showcase the versatility of this new chemo-optogenetic toolkit with respect to spatiotemporal control of gene expression in *E. coli*. On a structural level it is striking that most insertion sites resulting in switchable AraC behavior are located within the region between the ligand-binding domain (LBD) and the DNA-binding domain (LBD) of AraC (Fig. 4e). This trend can be explained by the functional role this region has, serving as dimerization interface upon AraC activation and by mediating the relative flexibility of both domains^32, 33^. It is thus no surprise that LOV domain insertions in this area can influence AraC function. Of note, AF2 structure predictions of AraC-I113-LOV and AraC-S170-LOV capture the former variant in a more compact conformation, which is in agreement with the less flexible repressor state of wildtype AraC^32, 33^ (note: AF2 predicts the LOV2 structure in its dark-adapted state) (Extended Data Fig. 7). AraC-S170-LOV, in turn, was predicted to have a more relaxed conformation, as would be expected for an active AraC (Extended Data Fig. 7). To further investigate the robustness of allosteric coupling in both hybrid proteins, we screened a set of point mutants for their effects on wildtype AraC and its engineered derivatives (Extended Data Fig. 8). The majority of mutations did not improve the AraC-I113-LOV switch, but rather reduced reporter activity, in the active (light) state or increased leakiness, i.e. reporter activity in the dark. Excitingly, several AraC-S170-LOV point mutants (e.g. T50S, G141D and V284F; mutations correspond to residues in wildtype AraC) showed an increased level of activity in the dark as compared to the initial variant, while likewise retaining potent reporter repression upon illumination. The mutations E3I and T241C, in turn, permanently impaired the function of the AraC-S170-LOV variant, while having no significant effect on AraC-I113-LOV. Finally, none of the tested mutations had major effects on the activity of wildtype AraC. Collectively, our data highlight (i) the variant-specificity of mutational effects in the engineered allosteric AraC-LOV hybrids and (ii) their increased functional and likely structural sensitivity towards minor sequence alterations. Moreover, the mutational data in conjunction with the arabinose-dependency data (Fig. 4b) indicate the interconnection of the natural arabinose-mediated allosteric regulation with a LOV-induced artificial allosteric pathway.

## Discussion

In this study, we investigated constraints of domain insertion engineering at the functional and structural level. Thereby, we considerably extended the existing body of work towards new protein families and, for the first time, compared the insertion tolerance of several evolutionary unrelated proteins side-by-side using directly effector protein function as readout. In agreement with previous studies^16, 17^, our data showcases the absence of any simplistic explanations for domain insertion permissibility. In contrast, we demonstrated that gradient boosting classifiers can help to decipher the importance of factors underlying domain insertion tolerance. Our models identified MSA-derived conservation statistics as main determinants of domain insertion tolerance, thus suggesting an evolutionarily informed approach to be particularly promising for domain insertion engineering (Fig. 3). In this context, parallels to statistical coupling analysis (SCA) and the idea of protein sectors can be drawn^5, 13^. While SCA analyses co-evolution of residues on a purely mathematical basis, however, our work underlines the indicative value of evolutionary insertion/deletion events. We note that in context of domain insertions, the predictive power of machine learning models is still constraint by the amount of available training data, which is, in turn, restricted by the current experimental capacity limits. The use of experimental data, such as the presented insertion library screens, in combination with larger datasets extracted from public protein sequence databases might provide an elegant solution to address this limitation in data size.

With respect to allosteric proteins, the screening pipeline developed here was efficient in identifying allosteric switches (Fig. 4, Extended Data Fig. 6). In previous work, a GFP-maltose-binding protein insertion library was enriched alternatingly in presence and absence of the input trigger in three consecutive rounds^12^. Our adaption of the method using parallel enrichment of the same library under different conditions (here culturing samples in presence or absence of light) turned out to be sufficient to reliably identify light-switchable proteins. A more stringent selection regime during FACS could potentially render even a single round of enrichment sufficient, which would further simplify and streamline the workflow for engineering of switchable effector proteins.

We note that several optogenetic bacterial expression systems exist^34–36^. These include the light-responsive AraC variant BLADE, which is based on the Vivid LOV domain from *Neurospora crassa* functioning via light-induced AraC dimerization^34^. In contrast to these previous examples, the transcription factors developed here are co-dependent on two stimuli, namely light and arabinose. This has interesting implications for synthetic biology applications and gene circuit control. Transcription factors co-dependent on two inputs enable the independent control of the state (on/off) and amplitude of activation for genetic programs. Previously, the combination of chemically inducible transcription factors and light-responsive regulators had to be combined within far more complex circuits to achieve the same goal^36–38^. The optogenetic variants presented here highly simplify such experimental setups by reducing the underlying system to a single protein component (see Supplementary note 2). Such single-protein Boolean logic gates could considerably streamline the design and increase the robustness of complex genetic circuits and biocomputing programs by reducing the number of required components and through the direct integration of signals within a single molecule.

In summary, our study pinpoints determinants of domain insertion tolerance and showcases the power of unbiased domain insertion screens for the engineering allosteric effector proteins with applications in synthetic biology and beyond.

## Methods

### Molecular cloning

All constructs used in this study are listed in Supplementary Table 1. The corresponding amino acid sequences of the encoded proteins are shown in Supplementary Table 2. Plasmids were constructed using Golden Gate assembly^39^. In brief, DNA fragments were amplified by PCR (Q5 2x Master Mix, New England Biolabs (NEB)), with primers carrying type IIS restriction enzyme recognition sites in their 5’-overhangs, which enabled the scarless assembly of constructs. PCRs were performed according to the NEB standard protocols. For Golden Gate assembly, the procedure described by Engler et al. was followed^39^. DNA-oligonucleotides were ordered from Merck and Integrated DNA Technologies (IDT). Double-stranded DNA fragments were purchased at IDT. Point mutants were cloned by introducing the changes via mismatching primers upon amplification of the full plasmid and subsequent phosphorylation and ligation. PCR products were resolved on 0.5x Tris-acetate-EDTA (TAE) 1 % agarose gels and the corresponding bands were cut out and purified using the QIAquick Gel Extraction kit (Qiagen). Restriction enzymes and T4 DNA ligase were obtained from NEB and Thermo Fisher Scientific. Following DNA assembly, Top10 *E. coli* cells (Thermo Fisher Scientific) were transformed with the respective construct, plated on agar and incubated overnight at 37 °C. Liquid cultures were inoculated from single colonies and grown overnight at 37 °C while shaking at 220 rounds per minute (rpm). DNA was purified using the QIAamp DNA Mini kit (Qiagen). All constructs were sequence-verified using Sanger sequencing (Microsynth Seqlab and Genewiz). The plasmid pTKEI-Dest, which served as backbone for the insertion libraries, was a gift from David Savage (Addgene plasmid # 79784; http://n2t.net/addgene:79784; RRID:Addgene_79784)^12^.

### Reporter assays

All reporter circuits used the monomeric red fluorescent protein 1 (RFP) as readout^40^. The design of the genetic circuits is depicted in Extended Figure 1. In short, the AraC reporter was created by placing the RFP coding sequence under control of a pBAD promoter. In case of the Flp recombinase, RFP was expressed from a constitutive promoter (J23102, http://parts.igem.org/Promoters/Catalog/Anderson). However, the coding sequence was inverted and flanked by Flp recognition target (FRT) sites. In the ground state, a dysfunctional mRNA is transcribed and only upon inversion of the RFP open reading frame by the recombinase, RFP is expressed. To measure TVMV protease activity, a ssrA-like degradation tag^41^ was fused to a constitutively expressed RFP; a TVMV recognition site was placed in between RFP and the degradation tag. Active TVMV protease would thus cleave off the degron resulting in RFP stabilization and an increase in fluorescence. Many related potyvirus proteases undergo a process called autolysis^42^, during which the protease cleaves off its own C-terminal region albeit at low efficiency. This results in a truncated protease with decreased activity. To ensure that only one TVMV protein species would be present during all assays, a previously reported, truncated TVMV version^43^ was used for insertion library generation. Finally, a reporter for SigF was constructed, based on a SigF-specific promoter design previously reported by Bervoets et al.^44^.

### Domain insertion library generation

To generate insertion libraries covering all possible effector protein positions, we used saturated programmable insertion engineering (SPINE)^15^. In short, the protein of interest was subdivided into chunks of ∼50 amino acids. For each chunk, an oligonucleotide sub-pool (Agilent) was designed, comprising 50 individual DNA sequences, each of which carried a Type IIS restriction enzyme recognition site handles behind a specific amino acid encoding triplet. A python pipeline for the automatic design of the required DNA sequences provided by Coyote-Maestas et al.^15^ was employed for oligo pool design. The sub-pools were then individually cloned into an expression vector carrying the full-length coding sequence of the respective effector protein of interest and transformed into chemically competent Oneshot Top10 *E. coli*. To ensure at least 40-fold coverage of the library, serial dilutions were plated on agar plates following transformation and the number of colony-forming units was calculated. The plasmid sub-libraries were purified from the bacteria using the QIAamp DNA Mini Preparation Kit (Qiagen). The DNA concentration was measured using the Quant-iT dsDNA (HS) assay kit (Thermo Fisher Scientific) and all sub-libraries for each individual effector protein were pooled using equal DNA concentrations. To ensure that no wildtype protein contamination was carried on during cloning, the insertion handle was replaced by a kanamycin expression cassette via Golden Gate assembly. *E. coli* cells were transformed and plated on three 20 cm LB-agar plates, supplemented with 50 µg/ml chloramphenicol and 25 µg/ml of kanamycin (Carl-Roth). Again, a library coverage of at least 20× was ensured by serial dilutions and colony counting. The next day, each plate was rinsed with 3 ml of LB and the colonies were gently scraped off with a spatula. The resulting liquid cultures were collected from the plates and pooled for each protein. Plasmid DNA was then purified from the cultures and the kanamycin handle was replaced by the insert domain of choice, again using Golden Gate cloning. Finally, Oneshot Top10 *E. coli* carrying the respective reporter plasmid were transformed with the assembled libraries by electroporation. Following recovery in super optimal broth supplemented with 20 mM glucose (Carl Roth) (SOC) for one hour at 37 °C and 220 rpm, transformed cells were grown in LB (50 µg/ml chloramphenicol and 25 µg/ml of kanamycin) overnight. Serial dilutions plated on agar were performed. Plates were incubated overnight, and a library coverage was estimated from colony counts (coverage was >50-fold for all samples). Finally, glycerol stocks of the libraries were prepared, by mixing the cultures with sterile 50 % (v/v) glycerol at a ratio of 1:1, and stocks were stored at −80 °C until usage.

### FACS-based library enrichment

Precultures of LB media (50 µg/ml of chloramphenicol and 25 µg/ml of kanamycin) were inoculated from glycerol stocks of *E. coli* strains carrying the insertion libraries. Positive control samples expressing the wildtype effector protein without insert, as well as negative controls expressing a different protein of similar size (not activating the reporter) from the same plasmid backbone, were included. The precultures were incubated for 16 h at 37 °C while shaking at 220 rpm. The next day, 1 ml LB cultures were inoculated with 10 µl from the precultures. These main cultures were supplemented with 16 mM L-arabinose and 400 µM IPTG for AraC, 400 µM IPTG for the TVMV protease, 200 µM IPTG for Flp, 100 µM IPTG for SigF for the first enrichment round and 200 µM for SigF during the second round of enrichment. These cultures were incubated for 16 h at 37 °C while shaking at 220 rpm. For the AraC-LOV2 libraries, two identical replicates were generated, one of which was incubated under blue light illumination and the other one in the dark. The next morning, the samples were diluted 1:100 in 1×PBS (Thermo Fisher Scientific) and kept on ice until sorting. FACS was performed on a FACSAria Fusion flow cytometer (BD Biosciences) at the ZMBH FACS facility (Heidelberg University). *E. coli* cells were identified and gated using the forward scatter (FSC) and side scatter (SSC) values (Supplementary Fig. 13). The red fluorescent peak was sorted from each library. If no clear peak was visible, the 5 % cells with the highest RFP levels were sorted. 25,000 cells were sorted for each library into LB media. Next, the collected cells were recovered for one hour in LB media without antibiotics at 37 °C and shaking at 220 rpm. Subsequently, 50 µg/ml chloramphenicol and 25 µg/ml of kanamycin were added, followed by incubation of cultures overnight. The next day, glycerol stocks were prepared from the cultures representing sorted libraries. A second round of FACS-sorting and enrichment was performed by repeating the procedure starting from the glycerol stocks after the first round of enrichment. FACS data was analyzed using the cytoflow python package (https://cytoflow.github.io/).

### Next generation sequencing

The input libraries, as well as the enriched sorted fractions were objected to heat lysis. Cells were pelleted and resuspended in water. Aliquots were heated to 95 °C for 10 min, followed by centrifugation at 10,000 g for 10 min to remove cell debris. The supernatant was transferred to new tubes and stored at - 20 °C until further use. The coding sequence of the libraries was amplified using the Q5 Hot Start High-Fidelity DNA Polymerase (NEB) and the PCR amplicons were separated from primer dimers on a 0.5x TAE 1 % agarose gel. The bands representing the protein hybrid libraries were excised and DNA was purified using the QIAquick Gel Extraction Kit (Qiagen). The DNA concentration was then measured with the Quant-iT dsDNA (HS) assay kit (Thermo Fisher Scientific) using a plate reader (Tecan Infinite 200 Pro). Next, the DNA was fragmented and the sequencing libraries were prepared using the Illumina Nextera XT kit (Illumina). The manufacturer’s protocol was followed, with two modifications. First, to prevent under-tagmentation, only 0.2 ng of DNA was used as input and the tagmentation step was performed for 15 min, instead of 5 min. Second, during library preparation, the samples to be pooled were barcoded using the Nextera XT Index Kit v2 (Illumina). The final sequencing libraries were then purified using AMPure XP magnetic beads (Beckman Coulter) according to the manufacturer’s protocol. A two-sided size selection was performed using 25 µl beads together with 50 µl input reaction during the first size selection step and 100 µl of beads during the second step. Following library clean-up, the DNA concentration was measured again using the Quant-iT dsDNA (HS) assay kit (Thermo Fisher Scientific) and the different libraries were pooled at equal concentrations. Next, library quality was assessed on a Bioanalyzer (Agilent) using the Agilent DNA 1000 Kit. Finally, samples were sequenced using the paired-end Illumina MiSeq and NextSeq sequencing services at the EMBL Gene Core facility (Heidelberg).

### Experimental characterization of individual variants from the domain insertion screen

Individual protein hybrids were isolated from the sorted fractions or cloned individually and stored as glycerol stocks in 25 % glycerol (Carl Roth). The variants tested are listed in Supplementary Table 1. Precultures of Oneshot Top10 cells carrying a RFP reporter plasmid specific to the respective protein hybrid, as well as a plasmid encoding the respective switchable variant, were inoculated from glycerol stocks into lysogeny broth (LB) (Carl Roth), supplemented with 50 µg/ml chloramphenicol (Carl Roth) and 25 µg/ml of kanamycin (Carl Roth). Cultures were prepared in technical triplicates in 96-well plates (Corning), using a volume of 200 µl per well. The precultures were incubated for 16 h at 37 °C while shaking at 220 rpm. Main cultures were similarly prepared in 96-well plates, using LB supplemented with 50 µg/ml chloramphenicol and 25 µg/ml of kanamycin, using the same induction scheme as for the FACS screen. The cultures were inoculated with 3 µl from the respective precultures and grown at 37 °C and 220 rpm for 16 h. Following incubation, RFP fluorescence and OD_600_ were measured on a plate reader (Tecan Infinite 200 Pro). For RFP measurements, an excitation wavelength of 490 nm and an emission wavelength of 520 nm were used. The reported RFP/OD600 values were calculated by dividing the measured fluorescence by the OD_600_ levels. Three independent biological replicates prepared and measured on different days were generated for each variant.

### Illumination setup

For the illumination of liquid cultures, custom-made LED setup was used. Eight blue light high-power LEDs (type CREE XP-E D5-15; emission peak ∼460 nm; emission angle ∼130°; LED-TECH.DE) were mounted onto an aluminum plate and connected to a Switching Mode Power Supply (Manson; HCS-3102). The LED-plate was installed upside down within a shaking incubator, so that the LEDs could illuminate the surface area of the shaking platform from a distance of approximately 30 cm. Liquid cultures were incubated in multi-well plates and illuminated at a constant intensity of 50 µmol/(m^2^·s).

For the illumination of agar plates (see “agar plate photography” below), a custom-made array of 96 LEDs (LB T64G-AACB-59-Z484-20-R33-Z, Osram, emission peak 469 nm, viewing angle 30 °, Mouser Electronics) mounted onto a circuit board was used, applying a light intensity of 15 µmol/(m^2^·s). This device was again powered by a Switching Mode Power Supply (Manson; HCS-3102). A photo-mask made from black vinyl (Starlab) was cut out by hand and was directly attached to the bottom of the agar plate. The plate was then placed above the LED array at a distance of ∼5 cm. The whole setup was installed inside a standard bacteria incubator (Minitron, Infors). The LED devices were custom-made by the workshop of the biology department at TU Darmstadt.

### Characterization of AraC-LOV2 hybrids

Precultures of Oneshot Top10 cells (Thermo Fisher Scientific) carrying the RFP reporter plasmid for AraC and an IPTG inducible expression plasmid encoding the transcription factor or its derivatives, were inoculated from glycerol stocks into LB (Carl Roth), supplemented with 50 µg/ml chloramphenicol (Carl Roth) and 25 µg/ml of kanamycin (Carl Roth). Cultures were prepared in 48-well plates (Corning), using a volume of 0.5 ml per well. The precultures were incubated for 16 h at 37 °C, while shaking at 220 rpm. Main cultures were similarly prepared in 48-well plates, using LB supplemented with 50 µg/ml chloramphenicol and 25 µg/ml of kanamycin, together with different amounts of IPTG (Carl Roth) and L-arabinose (Carl Roth). IPTG concentrations used in each sample are indicated in the corresponding figures/legends. The cultures were prepared in duplicates and inoculated with 5 µl from the respective precultures. Subsequently, one replicate was incubated under blue light exposure, while the other replicate was kept in the dark within the same incubator. The growth conditions were again 37 °C and 220 rpm for 16 h. Following incubation, RFP fluorescence and OD_600_ were measured in a plate reader. As before, an excitation wavelength of 490 nm and an emission wavelength of 520 nm were used and the fluorescence was normalized to the OD_600_. Experiments were performed in three independent replicates. Activity measurements of the AraC derivatives carrying point mutations were performed identically using an arabinose concentration of 8 mM.

### Agar plate photography

Prior to the experiment, agar plates were prepared using 1.5 % LB-agar, supplemented with 50 µg/ml chloramphenicol and 25 µg/ml of kanamycin, 400 µM IPTG and 25 mM L-arabinose (all Carl Roth). A preculture of the AraC-S170-LOV reporter strain was incubated overnight at 37 °C and 220 rpm. The next day, 0.6 % LB-agar was freshly prepared and cooled to ∼40 °C. Next, 3 ml of the liquid agar were supplemented with IPTG and L-arabinose to final concentrations of 400 µM and 25 mM, respectively. Finally, 300 µl of the preculture were quickly added to the agar, mixed by shaking and distributed on the previously prepared agar plates. After 30 minutes at room temperature, the top ager had solidified, and the photo-mask was glued to the bottom of the plate. Finally, the plate was incubated at 37 °C overnight, under constant blue light illumination. Images were acquired on the next day using a UV light source, high-pass filter and camera.

### Reversible optogenetic gene expression control

In a 48-well plate (Corning), 0.5 ml cultures were prepared, using LB media, supplemented with 50 µg/ml chloramphenicol and 25 µg/ml of kanamycin, 400 µM IPTG and 25 mM L-arabinose (all Carl Roth). The wells were inoculated with 5 µl of precultures that had been prepared as described above. The samples were then incubated at 37 °C and 220 rpm for three hours in darkness, followed by 3 h incubation under blue light exposure and a final step of 3 h in the dark. Prior to the first incubation step and after each following incubation period, the RFP fluorescence and the OD_600_ were measured in a plate reader. Following every incubation period the samples were diluted 1:30 into new plates with pre-warmed fresh media, containing all supplements. The final relative fluorescence was obtained by normalizing the RFP values to the measured OD_600_. Three independent replicates were generated by repeating experiments on different days.

### Structure prediction with AlphaFold2

Full-length structures of AraC, SigF, the TVMV protease, Flp, as well as the AraC-LOV2 fusions were obtained by AlphaFold2^21^ using the Colabfold implementation^22^. Structures were predicted using the “colabfold_batch” command with the “MMseqs2 (UniRef+Environmental)” MSA preferences. For the proteins without insertion, 5 models were run with three recycling iterations. To reduce compute time, only one model was predicted for the AraC-LOV2 hybrids, using a single recycling step. Images of the models were generated using UCSF ChimeraX (version 1.4)^45, 46^. To compute the position-wise RMSDs for between the AraC-LOV2 hybrids and the respective wildtype structures, the AF2 structures of AraC and the LOV2 domain were separately superimposed onto the prediction of the fusion proteins and RMSDs were calculated amino acid-wise. Computations were performed on the KIT Horeka cluster.

### NGS and data analysis

To analyze the sequencing data, fastq files were de-multiplexed using the Sabre tool (https://github.com/najoshi/sabre). The domain insertion frequencies were then calculated using a slightly modified version of the DIP-seq library^12^. Next the enrichment scores were determined using the following equation:

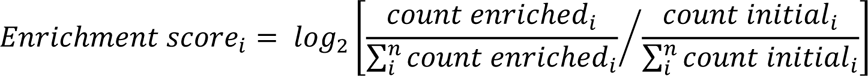

where *n* are the insertion positions within a given protein, *count enriched* represents the read counts after enrichment and *count initial* indicates the read counts of the initial library that was used as input to the sorting experiments. Insertions that were missing from the initial libraries were not taken into account during analysis. Insertion variants that entirely disappeared during sorting were assigned a log2 value of −10, which was in the range of the lowest obtained enrichment scores.

To gather position-wise protein features, diverse feature sources were used. Biophysical properties and linker propensity indices were fetched from the AAindex database^23, 24^. Information about secondary structure, accessible surface area and pLDDT score were extracted from the AF2-predicted structures. To map these features to the enrichment scores, the mean of the respective feature corresponding to the two amino acids that neighbor the insertion site were assigned to the enrichment. For the machine learning applications described below, the categorical features, such as secondary structures, were binarized similar to one-hot encodings, with the difference that every position could have two possible positive labels (if the secondary structure assignments of the two neighboring residues differ). The KLD, as well as the insertion and deletion statistics were based on sequence alignments. To this end, similar sequences were gathered using position-specific iterated basic local alignment search (PSI-BLAST)^47, 48^, with an expect threshold of 0.01 and a PSI-BLAST threshold of 0.005. The maximum number of sequences was limited to 5000. Based on these sequences, an MSA was calculated with MUSCLE^49^, using the Super5 algorithm with standard parameters. Finally, the KLD was calculated by the following equation:

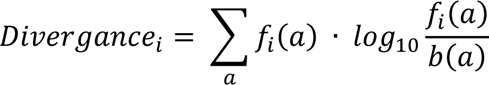

where the divergence is determined for the position *i* and *f(a)* is the frequency of the amino acid a at the given position, while *b(a)* represents the background frequency of the amino acid. Background frequencies were defined as the AA frequencies in SwissProt^50^. Of note, the definition of the gap background frequencies is non-trivial, as discussed by Teşileanu et al.^51^. Here, gaps were not included and the KLD is only based on AA frequencies. The position-wise insertion and deletion frequencies as well as the scores for the mean and median insertion lengths were calculated from pairwise alignments between the sequence of the protein of interest and its related sequences gathered by PSI-BLAST.

### Gradient boosting models

In order to train predictive models on the insertion data, the enrichment scores were first binarized. All sites exhibiting a positive enrichment were assigned the label 1 and all sites with negative insertions were labeled 0. All position-wise properties collected during data analysis were used as features. In addition, each amino acid and each secondary structure element represented individual additional features. Dataset construction and model training were performed using the Scikit-learn framework^52^. Individual datasets for every candidate protein, as well as a complete dataset using the combined data of all four proteins were constructed. A 80:20 train-test split was applied and the features were min-max scaled prior to training. Gradient boosting classifiers^29^ were trained using five-fold cross-validation. The hyperparameters were optimized on the complete dataset using grid search. For the final model, 100 estimators were trained using squared error and a learning rate of 0.1. The maximum depth of the trees was limited to four and the exponential loss was chosen. The maximum number of features parameter was kept at “auto”. The receiving operator characteristic and the average precision were chosen as performance metrics. The permutation importance and loss of impurity were calculated using the respective Scikit-learn functions.

## Data availability

Experimental raw data are available at Github under: https://github.com/Niopek-Lab/DI_screen. Plasmids encoding the AraC-I113-LOV, AraC-S170-LOV, AraC-S170-LOV_G141D and AraC-S170-LOV_T50S are available on Addgene (Addgene-IDs: xxxx, xxxx, xxxx, xxxx).

## Code availability

Jupyter notebooks of the computational analysis and additional scripts are available at Github under: https://github.com/Niopek-Lab/DI_screen.

## Acknowledgments

We thank the members of the Niopek lab for helpful discussions. Further, we are grateful to the ZMBH flow cytometry core facility (Heidelberg University) for support with cell sorting and the EMBL Genomics Core Facility (EMBL, Heidelberg) for performing deep sequencing. Finally, we sincerely thank the workshop at the Biology Department of the Technical University Darmstadt for the construction of customized illumination setups.

## Author contributions

D.N. and J.M. conceived the study. J.M., S.A. and P.B. designed and performed the experiments. J.M. implemented the computational analysis. D.N. directed the work and secured funding. J.M. and D.N. wrote the manuscript with support from all authors.

---

**Extended Data Fig. 1:**
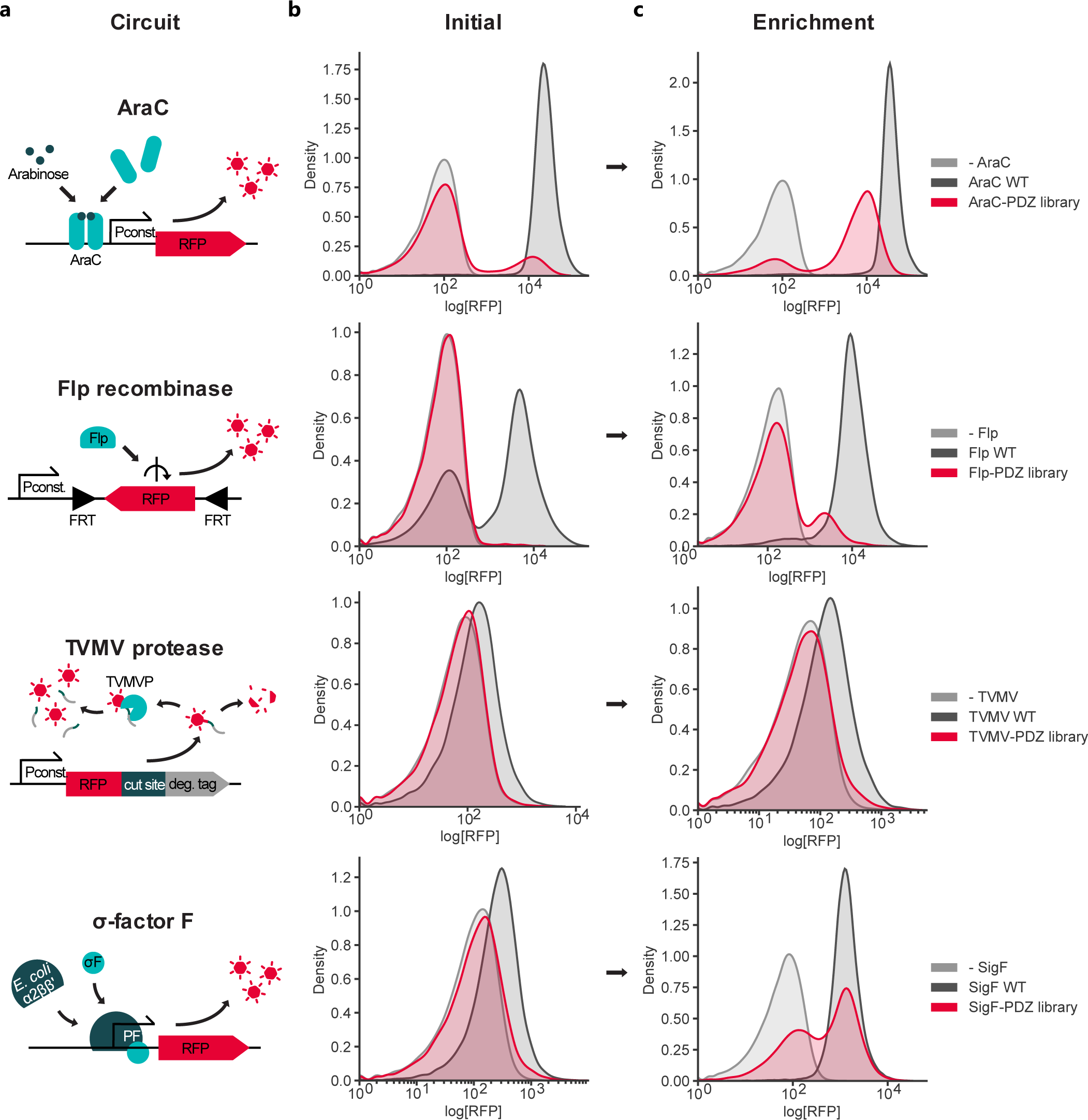
Enrichment of active effector-insert hybrids from candidate libraries via FACS. **a**, Schematics of the reporter assays for AraC, the Flp recombinase, the TVMV protease and SigF are shown. **b**, **c**, Histograms depicting the RFP fluorescence distribution during FACS-based library enrichment. Representative histograms generated from 25,000 gated events of (**b**) the initial library and (**c**) after the first enrichment are shown. The negative controls (-) carried a plasmid expressing a different candidate protein not activating the reporter construct.

**Extended Data Fig. 2:**
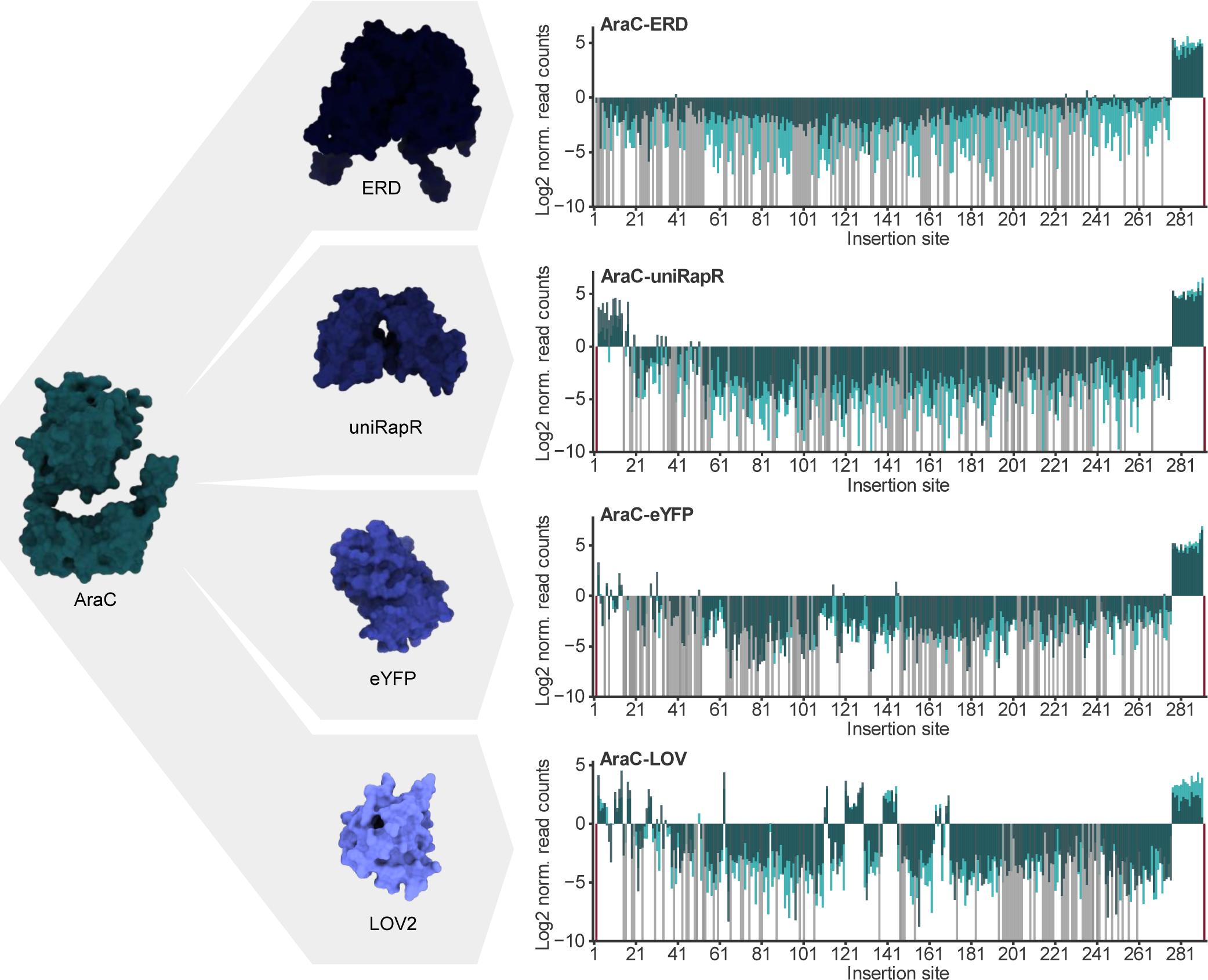
Domain insertion tolerance depends on the identity of the insert. Results from insertion screens of AraC with the ERD, LOV2, uniRapR and eYFP insert domains are shown. Enrichments are mapped to the respective insertion site as indicated by the position of the AraC preceding the insertion. Light green, dark green: individual replicates. Grey: variants with zero reads after enrichment. Red: variants missing in the initial library.

**Extended Data Fig. 3:**
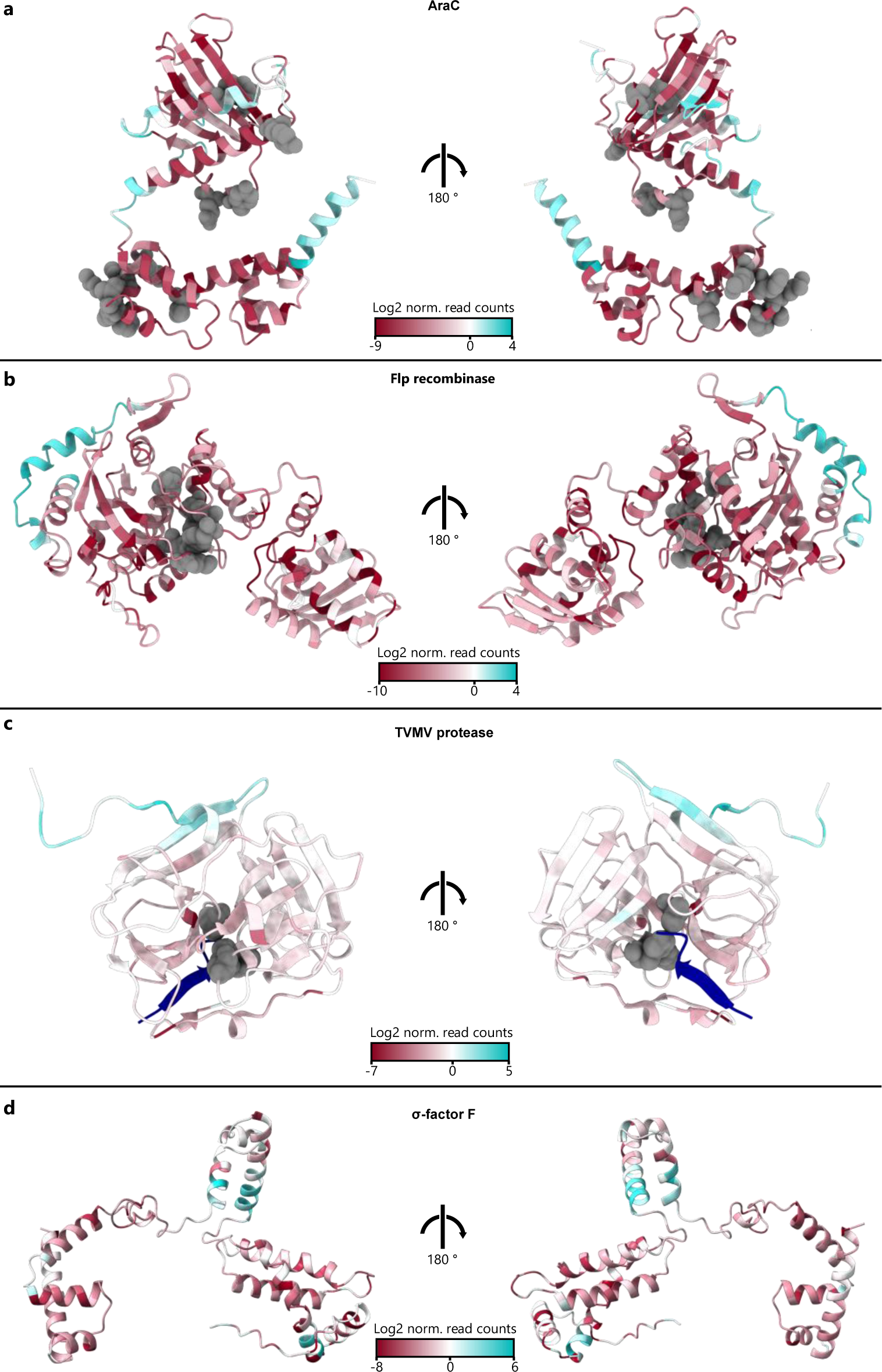
Positions with insertion tolerance are clustered at distinct, locally confined surface sites. **a-d**, The insertion scores from the PDZ libraries are mapped onto the AF2 structure predictions of AraC (**a**), and the Flp recombinase (**b**), a crystal structure of the TVMV protease (PDB-ID: 3MMG) (**c**) and an AF2 structure prediction of SigF (**d**). In **c**, the TVMV protease substrate is depicted in blue. Functionally critical residues are shown in grey for AraC, Flp and the TVMV protease.

**Extended Data Fig. 4:**
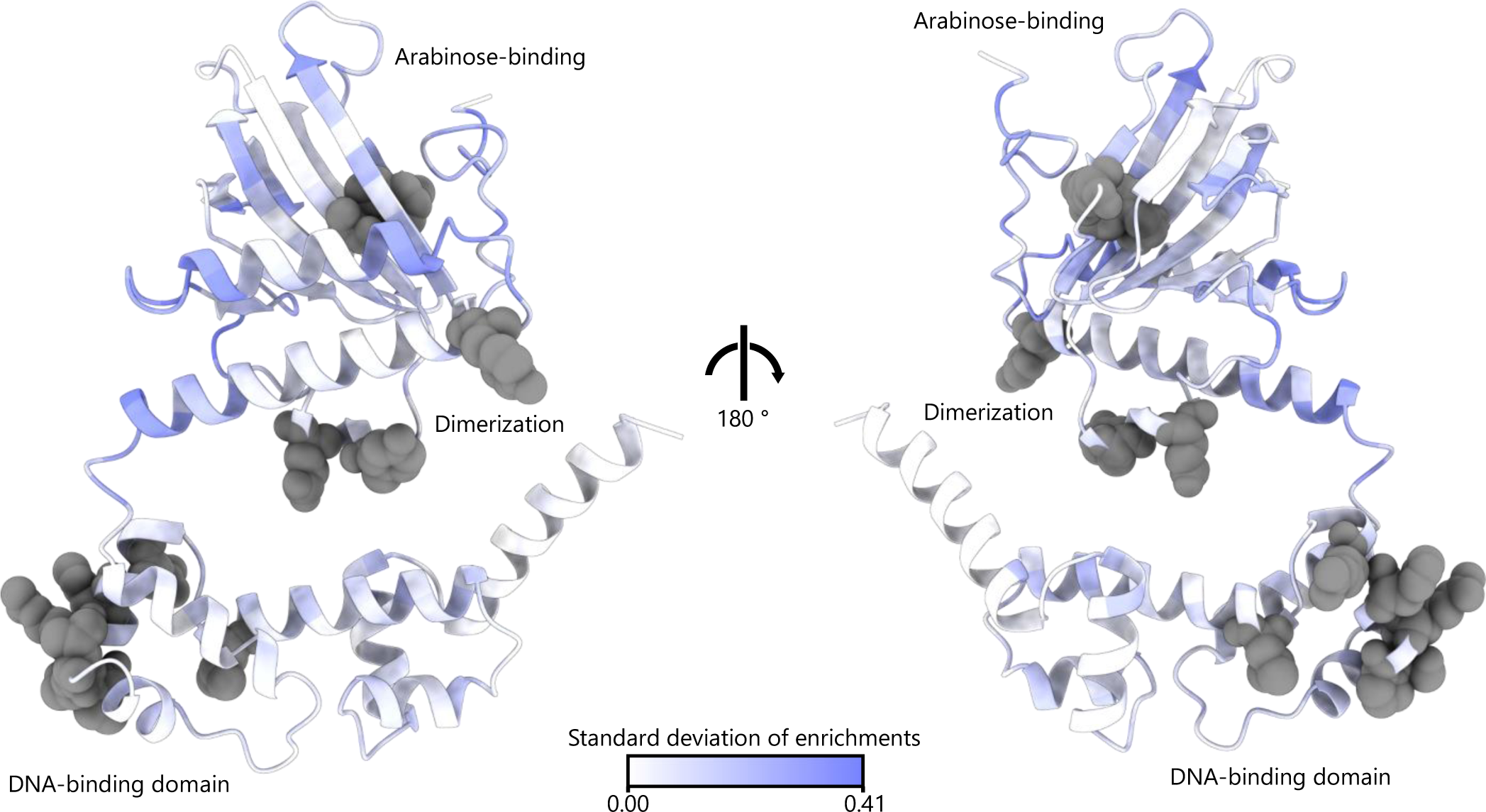
Insertion permissive regions are scattered across AraC and depend on the insert domain. The AF2-derived structure of AraC is colored by the SD of the min-max-scaled enrichment scores from all insert libraries corresponding to five different insertion domains. Functionally critical residues are highlighted in grey.

**Extended Data Fig. 5:**
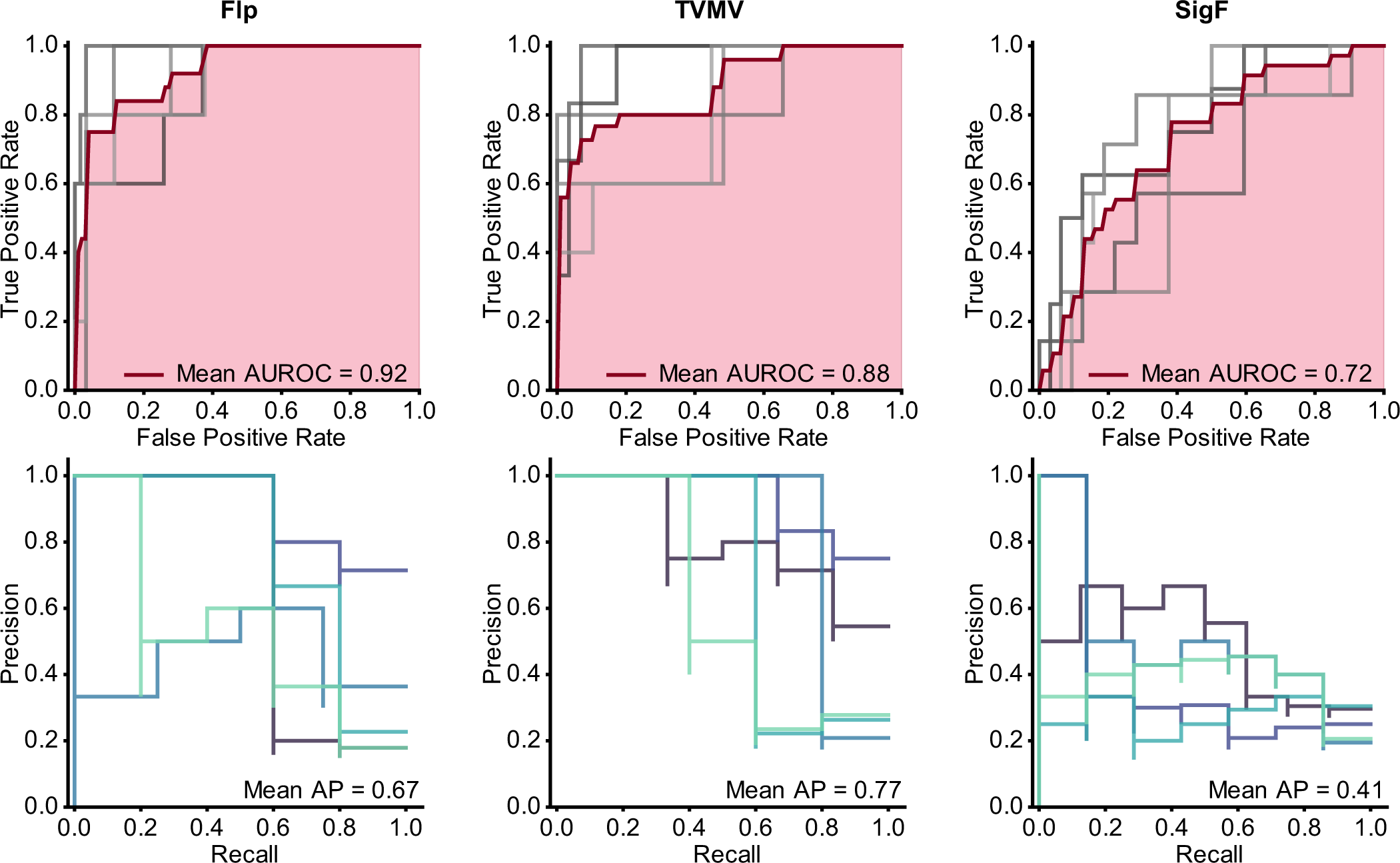
Gradient boosting models trained on positional features can infer insertion tolerance for individual proteins. Performance metrics of gradient boosting classifiers that were trained on the PDZ datasets for Flp, TVMV protease and SigF with five-fold cross-validation are shown. The ROC (top) and precision-recall curves (bottom) are depicted for individual folds. The mean ROC is shown in red and the mean AUC is marked in light red.

**Extended Data Fig. 6:**
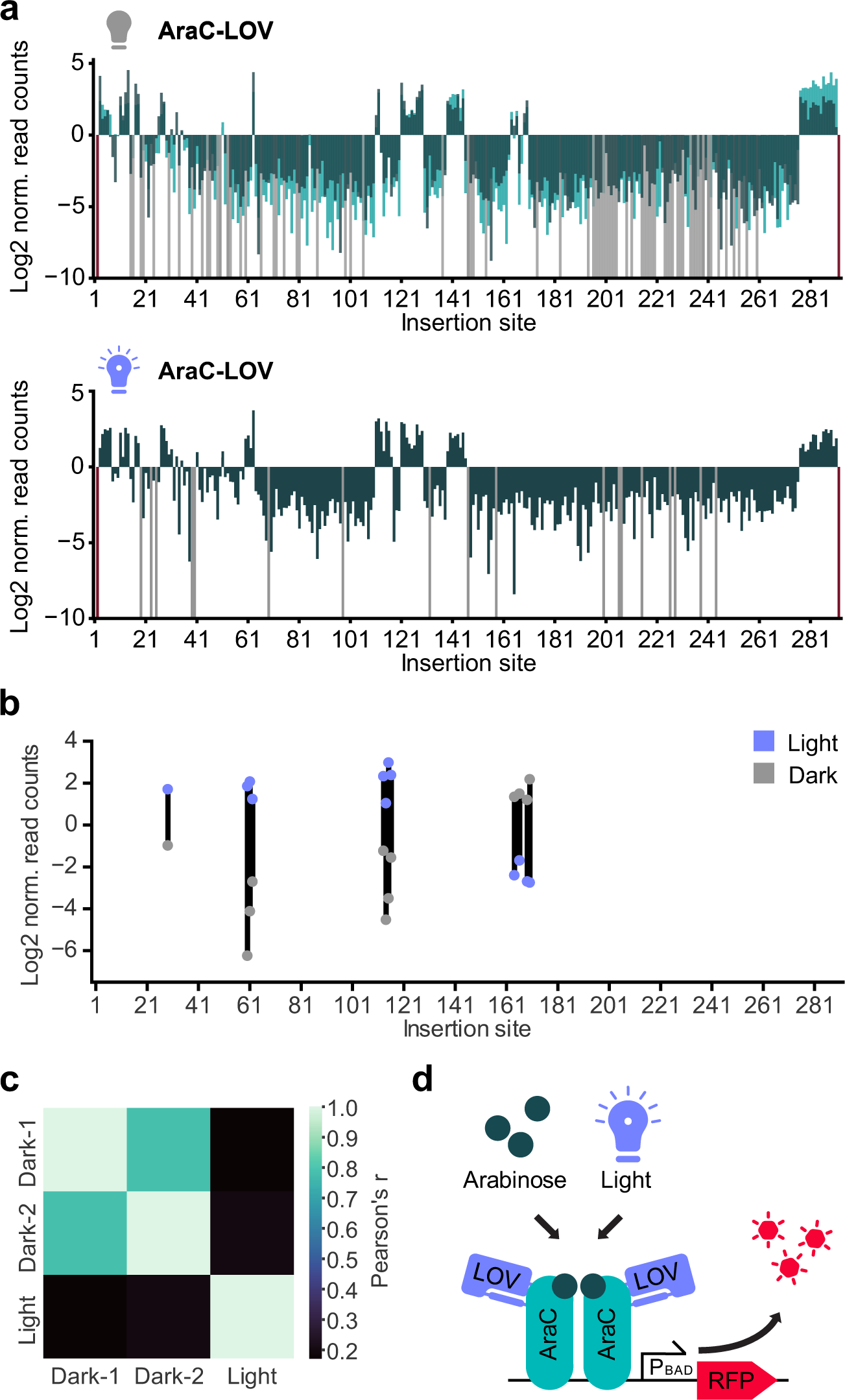
Distribution of light-switchable variants in the AraC-LOV2 dataset. **a**, Enrichment scores of AraC-LOV libraries that were sorted following incubation in darkness (upper panel) or under blue-light exposure (lower panel) are mapped onto the corresponding insertion sites of AraC (preceding the indicated residue). Values for the light exposed sample correspond to a single experiment. For the sample incubated in the dark, light green and dark green indicates individual replicates. Grey: variants with zero reads after enrichment. Red: variants missing in the initial library. **b**, Enrichment scores derived from experiments under light exposure or in darkness are marked by blue and grey points, respectively. Only datapoints from promising candidates with a log2 enrichment >1 in the active state and a log2 difference >2.5 between the light and dark states are shown. **c**, Pearson correlations between the different datasets are shown. Only positions of interest, that exhibited an enrichment in at least one replicate were included in the calculation. **d**, Schematic of the co-dependence of the AraC-LOV2 hybrids on arabinose and blue light.

**Extended Data Fig. 7:**
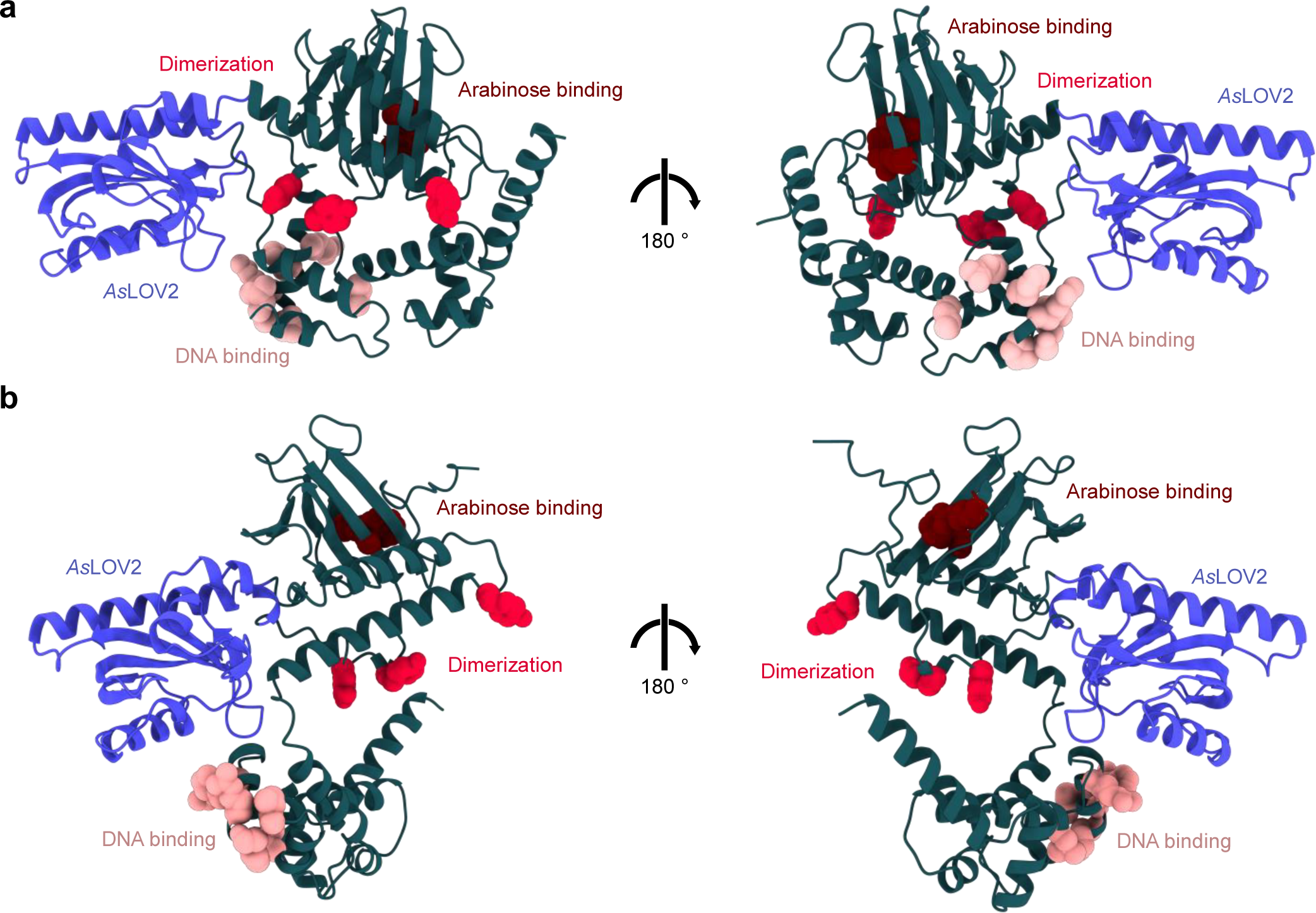
AlphaFold2 predicts different conformations for the lead AraC insertion variants. **a**, **b**, AF2 predictions of AraC-I113-LOV2 (**a**) and AraC-S170-LOV2 (**b**) are shown. AraC is depicted in green and the *As*LOV2 domain in blue. Residues that bind to the operator are highlighted in pink, key residues for dimerization in the induced state in red and the amino acids that are important for arabinose binding in vermilion.

**Extended Data Fig. 8:**
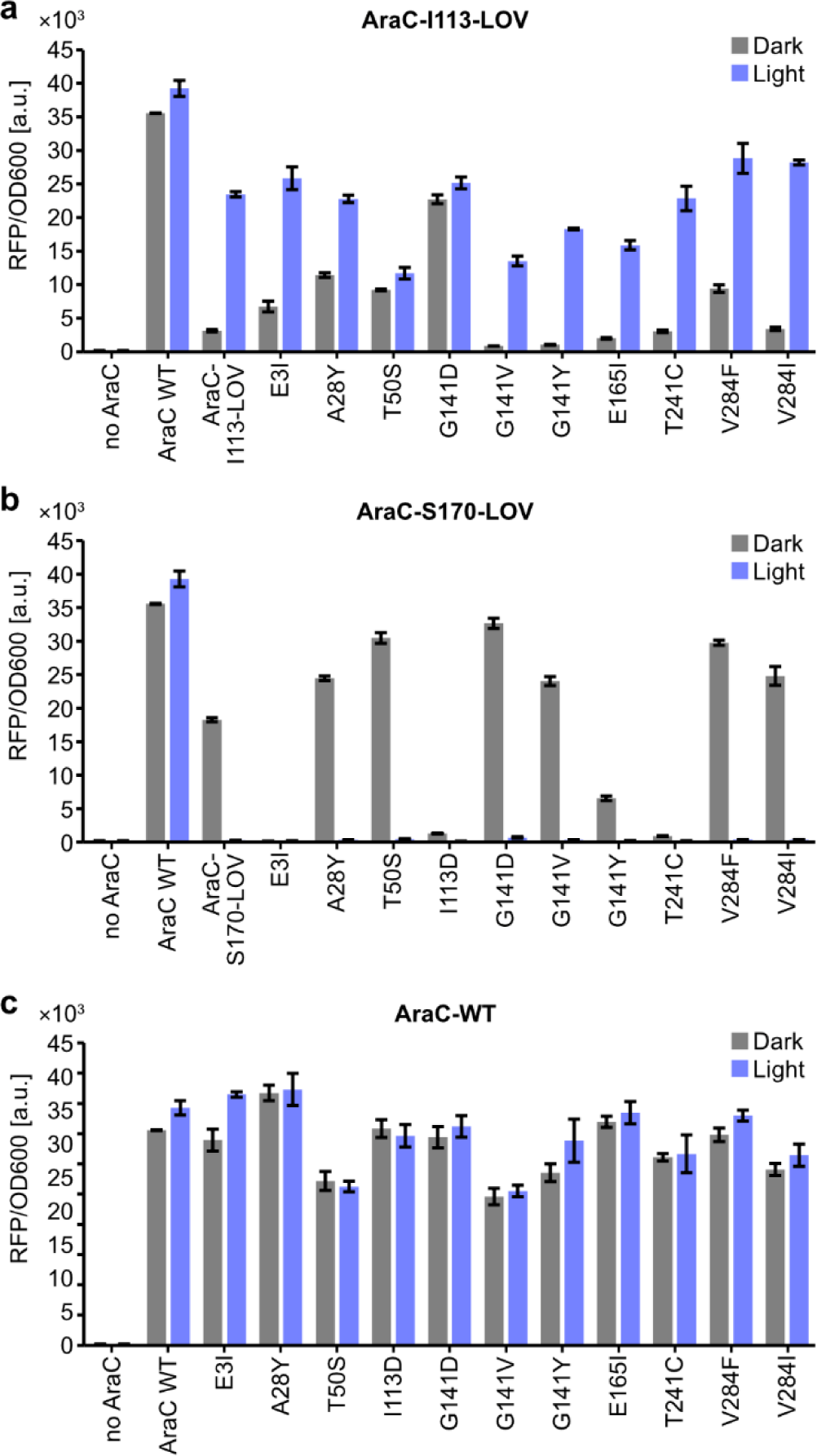
Point mutations improve the performance of the AraC-S170-LOV light switch. **a-c**, Cultures were inoculated from precultures carrying plasmids encoding an RFP reporter and the indicated AraC-I113-LOV (**a**), AraC-S170-LOV (**b**) and AraC (**c**) point mutants. The samples were incubated for 16 h under light exposure or in darkness at an arabinose concentration of 8 mM, followed by plate reader measurements of RFP fluorescence and OD600. Bars represent means from three independent biological replicates. Error bars show the SD.

**Extended Data Table 1:** Position-specific properties used for the analysis of domain insertion tolerance.

| Property | Description |
| --- | --- |
| ASA | Average surface accessibility |
| pLDDT | Position-wise pLDDT of AF2 models |
| Linker idx. Suyama | Linker propensity index<sup>25</sup> |
| Linker idx. George | Linker propensity index<sup>26</sup> |
| Linker idx. Bae | Linker index<sup>27</sup> |
| Hydrophobicity | Hydrophobicity<sup>53</sup> |
| Flexibility idx. | Flexibility index<sup>54</sup> |
| Molecular weight | Molecular amino acid weight |
| Average volume | Average amino acid volume |
| Positive charge | Positive charged amino acid |
| Negative charge | Negative charged amino acid |
| Net charge | Net charge of amino acid |
| Radius of gyration | Radius of gyration of the side chain |
| Side-chain stab. Idx. | Side-chain contribution to protein stability (KJ/mol)<sup>55</sup> |
| Buriability | Buriability<sup>56</sup> |
| KLD | KLD Kullback-Leibler divergence calculated from MSA |
| Insertion frequency | Insertion frequency in related natural sequences at the respective position |
| Deletion frequency | Deletion frequency in related natural sequences at the respective position |
| Mean ins. len. | Mean insertion length in related natural sequences at the respective position |
| Median ins. len. | Median insertion length in related natural sequences at the respective position |

## References

1. Ponting, C. P. & Russell, R. R. The Natural History of Protein Domains. Annu. Rev. Biophys. Biomol. Struct. 31, 45–71 (2002).

2. Jin, J. et al. Eukaryotic Protein Domains as Functional Units of Cellular Evolution. Sci. Signal. 2, (2009).

3. Dagliyan, O. et al. Engineering extrinsic disorder to control protein activity in living cells. Science 354, 1441–1444 (2016).

4. Siegel, M. S. & Isacoff, E. Y. A Genetically Encoded Optical Probe of Membrane Voltage. Neuron 19, 735–741 (1997).

5. Lee, J. et al. Surface Sites for Engineering Allosteric Control in Proteins. Science 322, 438–442 (2008).

6. Ostermeier, M. Engineering allosteric protein switches by domain insertion. *Protein Engineering*, Design and Selection 18, 359–364 (2005).

7. Guntas, G., Mansell, T. J., Kim, J. R. & Ostermeier, M. Directed evolution of protein switches and their application to the creation of ligand-binding proteins. Proceedings of the National Academy of Sciences 102, 11224–11229 (2005).

8. Dagliyan, O., Dokholyan, N. V. & Hahn, K. M. Engineering proteins for allosteric control by light or ligands. Nature Protocols 14, 1–21 (2019)

9. Hoffmann, M. D. et al. Optogenetic control of Neisseria meningitidis Cas9 genome editing using an engineered, light-switchable anti-CRISPR protein. Nucleic Acids Research 49, 1–11 (2021).

10. Bubeck, F. et al. Engineered anti-CRISPR proteins for optogenetic control of CRISPR–Cas9. Nature Methods 15, 924–927 (2018).

11. Oakes, B. L. et al. Profiling of engineering hotspots identifies an allosteric CRISPR-Cas9 switch. Nature Biotechnology 34, 646–651 (2016).

12. Nadler, D. C., Morgan, S. A., Flamholz, A., Kortright, K. E. & Savage, D. F. Rapid construction of metabolite biosensors using domain-insertion profiling. Nature Communications 7, 1–11 (2016).

13. Reynolds, K. A., McLaughlin, R. N. & Ranganathan, R. Hot spots for allosteric regulation on protein surfaces. Cell 147, 1564–1575 (2011).

14. Edwards, W. R., Busse, K., Allemann, R. K. & Jones, D. D. Linking the functions of unrelated proteins using a novel directed evolution domain insertion method. Nucleic Acids Research 36, e78 (2008).

15. Coyote-maestas, W., Nedrud, D., Okorafor, S., He, Y. & Schmidt, D. Targeted insertional mutagenesis libraries for deep domain insertion profiling. Nucleic Acids Research 48, e11 (2019).

16. Coyote-Maestas, W., He, Y., Myers, C. L. & Schmidt, D. Domain insertion permissibility-guided engineering of allostery in ion channels. Nature Communications 10, 290 (2019).

17. Coyote-Maestas, W. et al. Probing ion channel functional architecture and domain recombination compatibility by massively parallel domain insertion profiling. Nat Commun 12, 7114 (2021).

18. Fernandez-Rodriguez, J. & Voigt, C. A. Post-translational control of genetic circuits using Potyvirus proteases. Nucleic Acids Research 44, 6493–6502 (2016).

19. Ormö, M. et al. Crystal Structure of the Aequorea victoria Green Fluorescent Protein. Science 273, 1392–1395 (1996).

20. Dagliyan, O. et al. Rational design of a ligand-controlled protein conformational switch. Proceedings of the National Academy of Sciences 110, 6800–6804 (2013).

21. Jumper, J. et al. Highly accurate protein structure prediction with AlphaFold. Nature 596, 583–589 (2021).

22. Mirdita, M. et al. ColabFold: Making Protein folding accessible to all. Nature Methods 19, 679–682, (2022).

23. Kawashima, S. et al. AAindex: amino acid index database, progress report 2008. Nucleic Acids Res 36, D202–D205 (2008).

24. Kawashima, S. & Kanehisa, M. AAindex: Amino Acid index database. Nucleic Acids Research 28, 374 (2000).

25. Suyama, M. & Ohara, O. DomCut: prediction of inter-domain linker regions in amino acid sequences. Bioinformatics 19, 673–674 (2003).

26. George, R. A. & Heringa, J. An analysis of protein domain linkers: their classification and role in protein folding. *Protein Engineering*, Design and Selection 15, 871–879 (2002).

27. Bae, K., Mallick, B. K. & Elsik, C. G. Prediction of protein interdomain linker regions by a hidden Markov model. Bioinformatics 21, 2264–2270 (2005).

28. Akdel, M. et al. A structural biology community assessment of AlphaFold2 applications. Nat Struct Mol Biol 29, 1056–1067 (2022).

29. Friedman, J. H. Stochastic gradient boosting. Computational Statistics & Data Analysis 38, 367–378 (2002).

30. Louppe, G. Understanding Random Forests: From Theory to Practice. Preprint at https://doi.org/10.48550/arXiv.1407.7502 (2015).

31. Mathony, J. & Niopek, D. Enlightening Allostery: Designing Switchable Proteins by Photoreceptor Fusion. Advanced Biology 5, 2000181 (2021).

32. Soisson, S. M., MacDougall-Shackleton, B., Schleif, R. & Wolberger, C. Structural Basis for Ligand-Regulated Oligomerization of AraC. Science 276, 421–425 (1997).

33. Schleif, R. AraC protein, regulation of the l-arabinose operon in Escherichia coli, and the light switch mechanism of AraC action. FEMS Microbiol Rev 34, 779–796 (2010).

34. Romano, E. et al. Engineering AraC to make it responsive to light instead of arabinose. Nat Chem Biol 17, 817–827 (2021).

35. Dietler, J. et al. A Light-Oxygen-Voltage Receptor Integrates Light and Temperature. Journal of Molecular Biology 433, 167107 (2021).

36. Li, X. et al. A single-component light sensor system allows highly tunable and direct activation of gene expression in bacterial cells. Nucleic Acids Research 48, e33 (2020).

37. Jayaraman, P. et al. Blue light-mediated transcriptional activation and repression of gene expression in bacteria. Nucleic Acids Research 44, 6994–7005 (2016).

38. Jayaraman, P., Yeoh, J. W., Zhang, J. & Poh, C. L. Programming the Dynamic Control of Bacterial Gene Expression with a Chimeric Ligand- and Light-Based Promoter System. ACS synthetic biology 7, 2627–2639 (2018).

39. Engler, C., Kandzia, R. & Marillonnet, S. A One Pot, One Step, Precision Cloning Method with High Throughput Capability. PLOS ONE 3, e3647 (2008).

40. Campbell, R. E. et al. A monomeric red fluorescent protein. Proceedings of the National Academy of Sciences 99, 7877–7882 (2002).

41. McGinness, K. E., Baker, T. A. & Sauer, R. T. Engineering Controllable Protein Degradation. Molecular Cell 22, 701–707 (2006).

42. Kapust, R. B. et al. Tobacco etch virus protease: mechanism of autolysis and rational design of stable mutants with wild-type catalytic proficiency. *Protein Engineering*, Design and Selection 14, 993–1000 (2001).

43. Sun, P., Austin, B. P., Tözsér, J. & Waugh, D. S. Structural determinants of tobacco vein mottling virus protease substrate specificity: Structure of TVMV Protease/Substrate Complex. Protein Science 19, 2240–2251 (2010).

44. Bervoets, I. et al. A sigma factor toolbox for orthogonal gene expression in Escherichia coli. Nucleic Acids Research 46, 2133–2144 (2018).

45. Goddard, T. D. et al. UCSF ChimeraX: Meeting modern challenges in visualization and analysis. Protein Sci 27, 14–25 (2018).

46. Pettersen, E. F. et al. UCSF ChimeraX: Structure visualization for researchers, educators, and developers. Protein Sci 30, 70–82 (2021).

47. Altschul, S. F. et al. Gapped BLAST and PSI-BLAST: a new generation of protein database search programs. Nucleic Acids Res 25, 3389–3402 (1997).

48. Altschul, S. F., Gish, W., Miller, W., Myers, E. W. & Lipman, D. J. Basic local alignment search tool. J Mol Biol 215, 403–410 (1990).

49. Edgar, R. C. MUSCLE: multiple sequence alignment with high accuracy and high throughput. Nucleic Acids Res 32, 1792–1797 (2004).

50. Bairoch, A. & Apweiler, R. The SWISS-PROT protein sequence database: its relevance to human molecular medical research. J Mol Med (Berl) 75, 312–316 (1997).

51. Teşileanu, T., Colwell, L. J. & Leibler, S. Protein Sectors: Statistical Coupling Analysis versus Conservation. PLoS Computational Biology 11, 1–20 (2015).

52. Pedregosa, F. et al. Scikit-learn: Machine Learning in Python. Journal of Machine Learning Research 12, 2825–2830 (2011).

53. Prabhakaran, M. The distribution of physical, chemical and conformational properties in signal and nascent peptides. Biochem J 269, 691–696 (1990).

54. Bhaskaran, R. & Ponnuswamy, P. k. Positional flexibilities of amino acid residues in globular proteins. International Journal of Peptide and Protein Research 32, 241–255 (1988).

55. Takano, K. & Yutani, K. A new scale for side-chain contribution to protein stability based on the empirical stability analysis of mutant proteins. Protein Engineering 14, 525–8, (2001).

56. Zhou, H. & Zhou, Y. Quantifying the effect of burial of amino acid residues on protein stability. Proteins: Structure, Function, and Bioinformatics 54, 315–322 (2004).

