## Supplementary Information for "Dissecting the Determinants of Domain Insertion Tolerance and Allostery in Proteins"

### Content:

**Supplementary Note 1** | Analyzing Alphafold2 structure predictions of domain insertion variants 2

**Supplementary Note 2** | Optogenetic AraC variants and single-protein Boolean logic gates 3

**Supplementary Fig. 1** | Cloning of domain insertion libraries via SPINE yields near-complete coverage of domain insertion positions. 4

**Supplementary Fig. 2** | Domain insertion profiling outcomes are highly reproducible. 5

**Supplementary Fig. 3** | Cross-validation of the domain insertion screen by experimental characterization of individual insertion variants. 6

**Supplementary Fig. 4** | Enrichment scores mapped onto structures of the Flp-holliday junction complex. 7

**Supplementary Fig. 5** | AlphaFold2 predictions accurately capture the structures of the candidate proteins. 8

**Supplementary Fig. 6** | Correlations between the enrichment scores and surface accessibility or secondary structures. 9

**Supplementary Fig. 7** | Successful domain insertion cannot be predicted from amino acid identity. 10

**Supplementary Fig. 8** | Heatmap of pairwise Spearman correlations between all selected positional features. 11

**Supplementary Fig. 9** | The position-specific pLDDT scores of wildtype AraC do not correlate with domain insertion susceptibility. 12

**Supplementary Fig. 10** | Correlations of structure predictions with domain insertion susceptibility. 13

**Supplementary Fig. 11** | Full comparison of the trained classifier to baseline predictors. 15

**Supplementary Fig. 12** | Alignment-derived statistics are key predictors of insertion tolerance. 16

**Supplementary Fig. 13** | Gating strategy 17

**Supplementary Table 1** | Constructs created and used in this study. 18

**Supplementary Table 2** | Amino acid sequences of the used proteins and insert domains. 21

**Supplementary Table 3** | PDB IDs of protein structures shown and used in this study. 22

Supplementary References 23

**Supplementary Note 1 | Analyzing Alphafold2 structure predictions of domain insertion variants**

In light of the recent advances in protein structure prediction, a logical question was if AF2 could guide the identification of promising domain fusions. We chose the pLDDT metric as a starting point, as it was previously shown to be correlated with flexible protein regions[1–3], and could hence serve as potential indicator for suitable domain insertion sites. Analysis of the pLDDT scores of individual amino acids from an AF2-derived structure of wildtype AraC revealed a trend towards lower pLDDT values at enriched sites, although the resulting correlation was very weak (Spearman's r of -0.26; Supplementary Fig. 9).

Next, we predicted AF2 structures of all possible PDZ insertions into AraC (Supplementary Fig. 10a). Representing all amino acid-wise pLDDT scores corresponding to AraC from each fusion protein in a heatmap allowed us to investigate the effect each insertion has on the pLDDT scores of AraC (Supplementary Fig. 10b). Most prominent in the resulting representation is a diagonal of decreased pLDDT values corresponding to the residues neighboring the respective position of the PDZ insertion. These lower values could implicate structural flexibility around the respective insertion site. The interpretation is backed by the fact that the unstructured loops of AraC are also visible as vertical regions with decreased pLDDT scores. We note that the structure of the N-terminal β-barrel (AA 20-100) is implicitly visible in the heatmap by a symmetric pattern of locally decreased pLDDT scores indicating its loop regions in the upper left quarter. Indeed, the pLDDT scores reflected structural features of AraC and potentially local conformational effects of insertions, albeit these findings remain speculative as this point. However, the pLDDT score changes did not correlate with the experimentally determined enrichment scores (Supplementary Fig. 10c).

In line with the pLDDT values, the structural differences between predicted models of wildtype AraC and the corresponding parts of AraC-PDZ hybrid structures exhibited a similar trend (Supplementary Fig. 10d). When, in turn, the PDZ insert was compared to its wildtype conformation (Supplementary Fig. 10e), misfolding of the domain was predicted for several hybrids, although the corresponding insertion sites did not necessarily correspond to regions of significant depletion in our screen. Taken together, the exploration of predicted hybrid protein structures suggested that AF2 is not able to capture the functional effects of domain insertions in a meaningful way. Given the generally lower performance of AF2 on multi-domain proteins[4], domain insertion engineering might still be beyond the scope of AF2 and similar state-of-the-art structure prediction methods. Nonetheless, AF2 predictions do reflect diverse structural features of AraC.

**Supplementary Note 2 | Optogenetic AraC variants and single-protein Boolean logic gates**

Boolean logic computations are typical elements of genetic circuits and programs used in synthetic biology. They are usually implemented at the transcriptional and/or translational level[5,6], which causes delays in the signal relay and integration. Protein-based logic computation in contrast does not suffer from these limitations and thus holds great potential for the custom control of cellular processes and the implementation of computational circuits in cells[7,8]. However, increasingly complex circuit designs require the use of a large number of individual protein components as well as their efficient communication via protein-protein interactions[7,8]. A recent preprint, for instance, impressively demonstrated design of neural-network computations on the protein level[9], showcasing the increasing power of artificial protein networks in living cells. Such complex cellular compute programs, however, show considerable noise and potentially cross-talk within the system and thus require efficient processing at the level of the individual protein components.

In contrast to such commonly used protein logic gate designs based on several separate protein components, our AraC-LOV2 fusions represent single protein Boolean logic gates (Fig. 4d). AraC-I113-LOV2 acts as an AND-gate, integrating blue light and arabinose as inputs, while AraC-S170-LOV2 represents a NIMPLY-gate. We have found only two other examples of engineered, single-protein logic gates in the literature. The first was constructed by fusing LOV2 and uniRapR domains to a kinase[10] resulting in an OR gate behavior, while the second comprises an engineered transcription factor that responds to light and temperature changes[11]. A particular advantage of these single protein-based logic gates is that they provide a direct wiring from the input signals to the output computation within a single polypeptide chain, a computation that would otherwise require several separate protein components. Building on this unique feature could highly simplify the design of compute circuits and their operation in living cells. Our combination of the naturally existing allosteric signaling in AraC with an artificial, second input might be an engineering approach that could be easily adapted to other proteins and thereby facilitate the implementation of Boolean logics for demanding computational operations in cells.

Taken together, the possibility to integrate complex information and wiring it to desired actuations is one of the main goals in synthetic biology. Integrating more functions into single amino acid chains therefore has the potential to simplify molecular networks, release metabolic burden from the host cell[12] and reduce noise derived from stochastic fluctuations of the individual components[13]. This way, single-protein logic gates could contribute to future generations of synthetic biology approaches to program and re-wire cells.

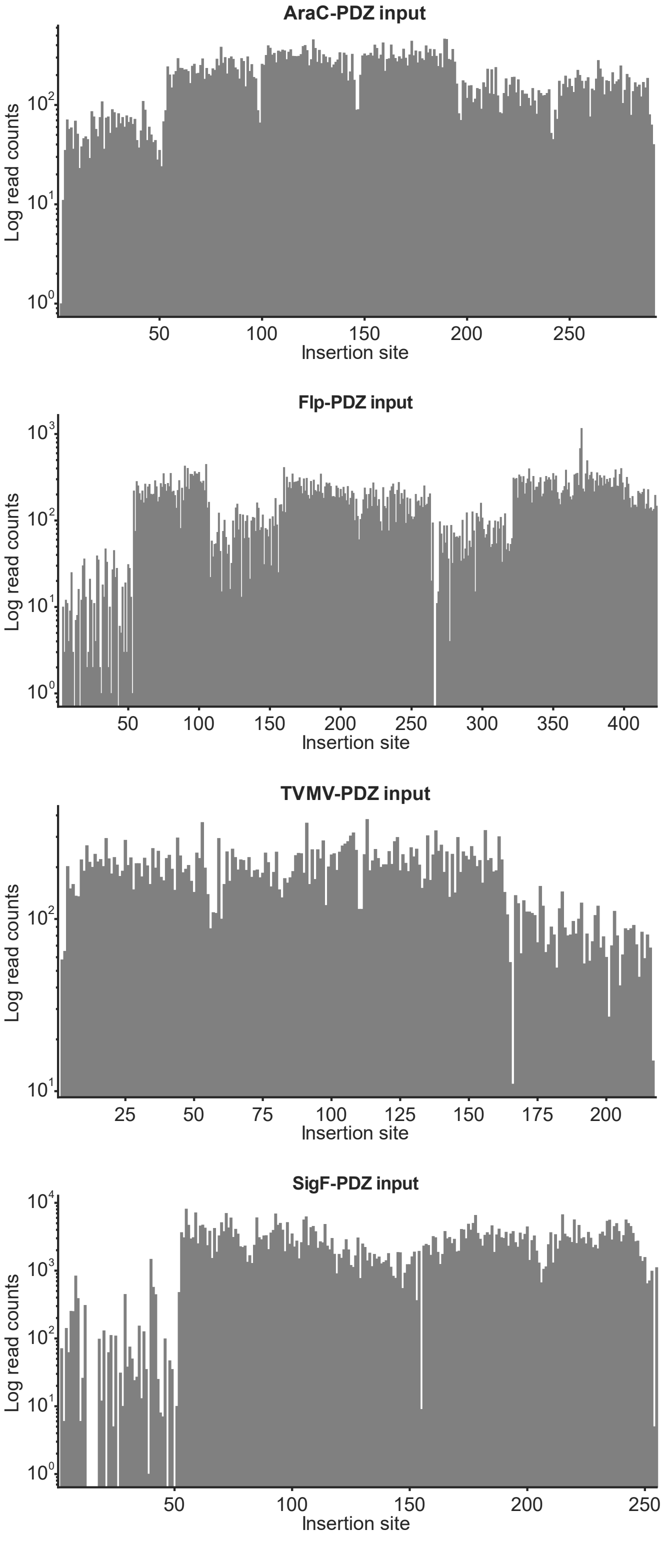


**Supplementary Fig. 1 | Cloning of domain insertion libraries via SPINE yields near-complete coverage of domain insertion positions.**

The insertion library coverage was assessed via NGS. Histograms represent the log-normalized read counts for insertions at the respective position (amino acid/codon).

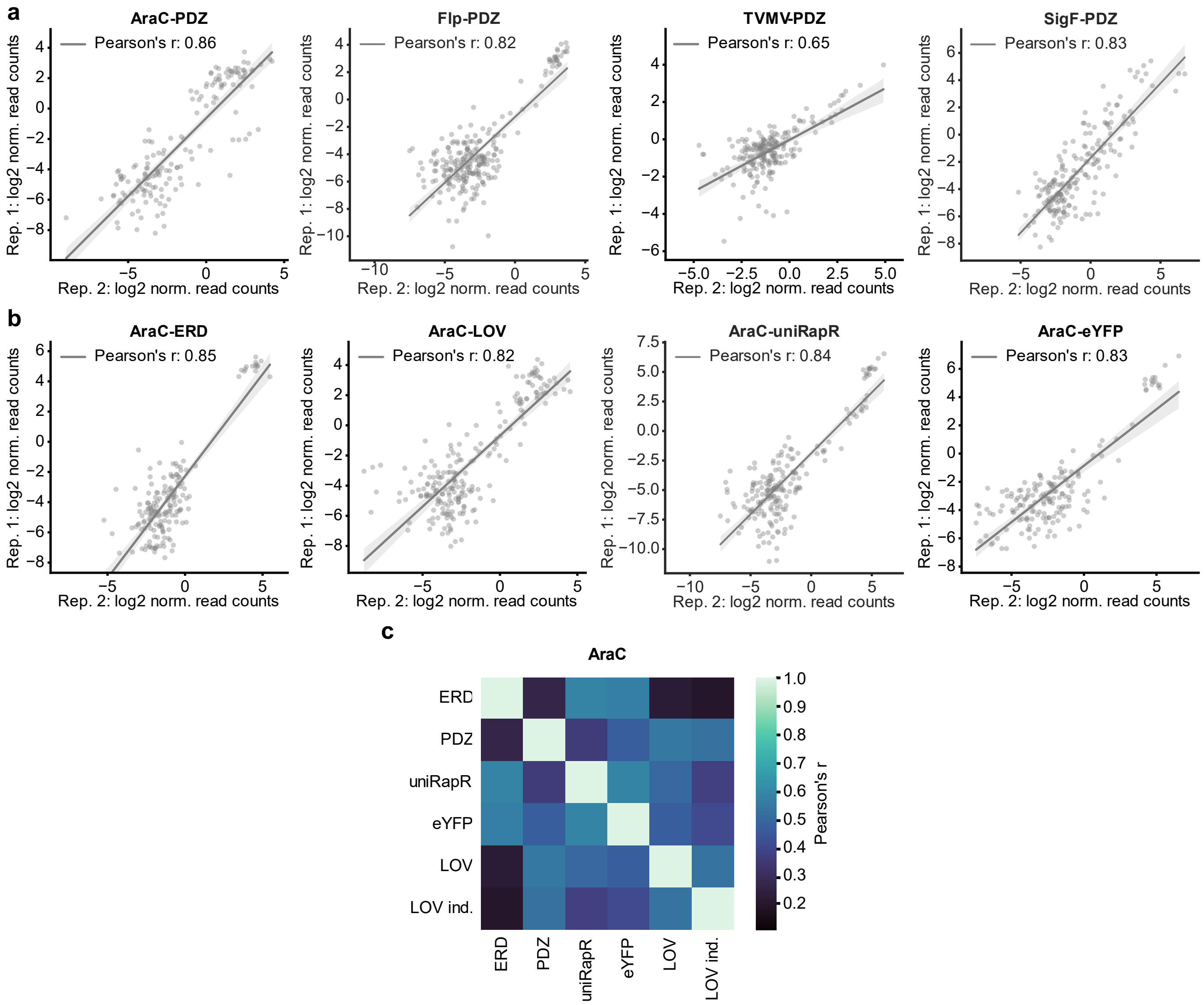


**Supplementary Fig. 2 | Domain insertion profiling outcomes are highly reproducible.**

**a-b**, The enrichment scores of biological replicate-1 are plotted against the respective scores from a second replicate-2 for the different effector-PDZ libraries (**a**) and the additional AraC libraries with varying insert domains (**b**). Only variants that were not fully depleted during enrichment are shown. A linear fit with 95 % confidence intervals is included and Pearson correlations coefficients are indicated. Rep., replicate; norm., normalized. **c**, The heatmap shows pairwise Pearson correlations between all domain inserted into AraC. Enrichments of the AraC-LOV2 library in darkness and under light induction (ind.) were assessed and are depicted separately.

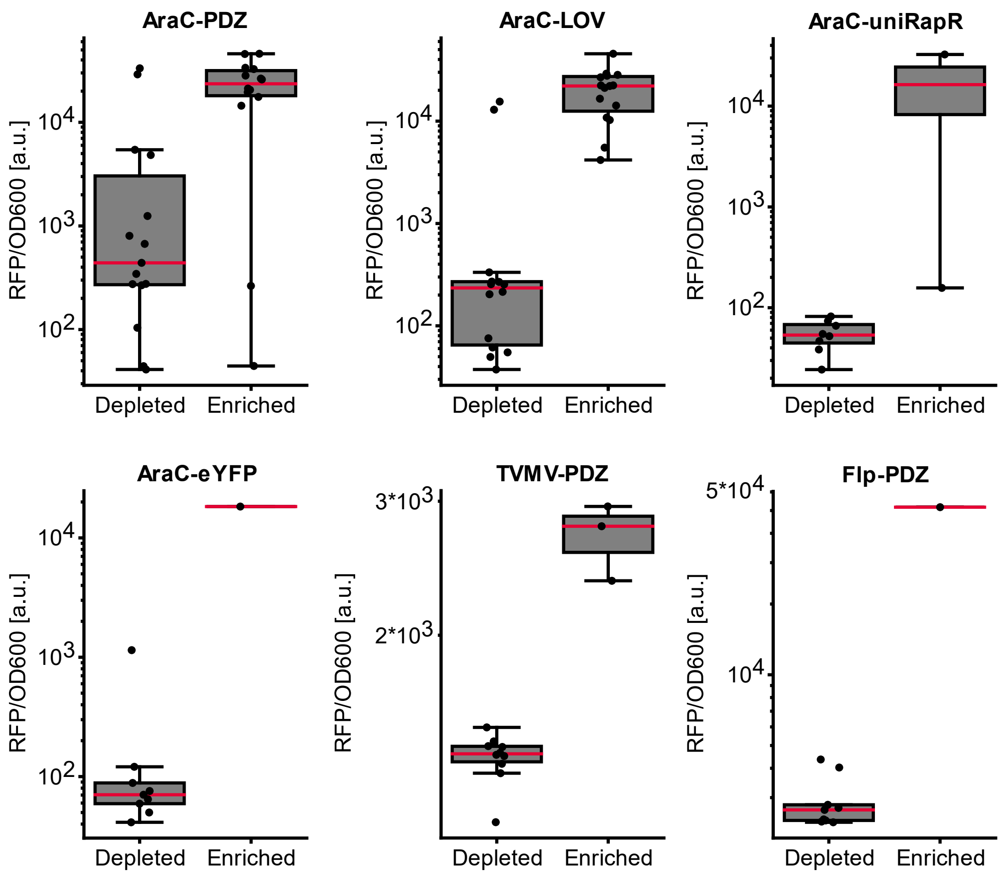


**Supplementary Fig. 3 | Cross-validation of the domain insertion screen by experimental characterization of individual insertion variants.**

Individual domain insertion variants were cloned and their activity was assessed using the respective RFP reporter assays. Boxplots indicate the resulting normalized fluorescence for enriched and depleted candidate. Individual data points correspond to the mean of three biological replicates, each of which reflect of three underlying technical replicates. The IQR is marked by the box and the median is represented by a red line. Whiskers extend to the 1.5-fold IQR or to the value of the smallest or largest enrichment, respectively.

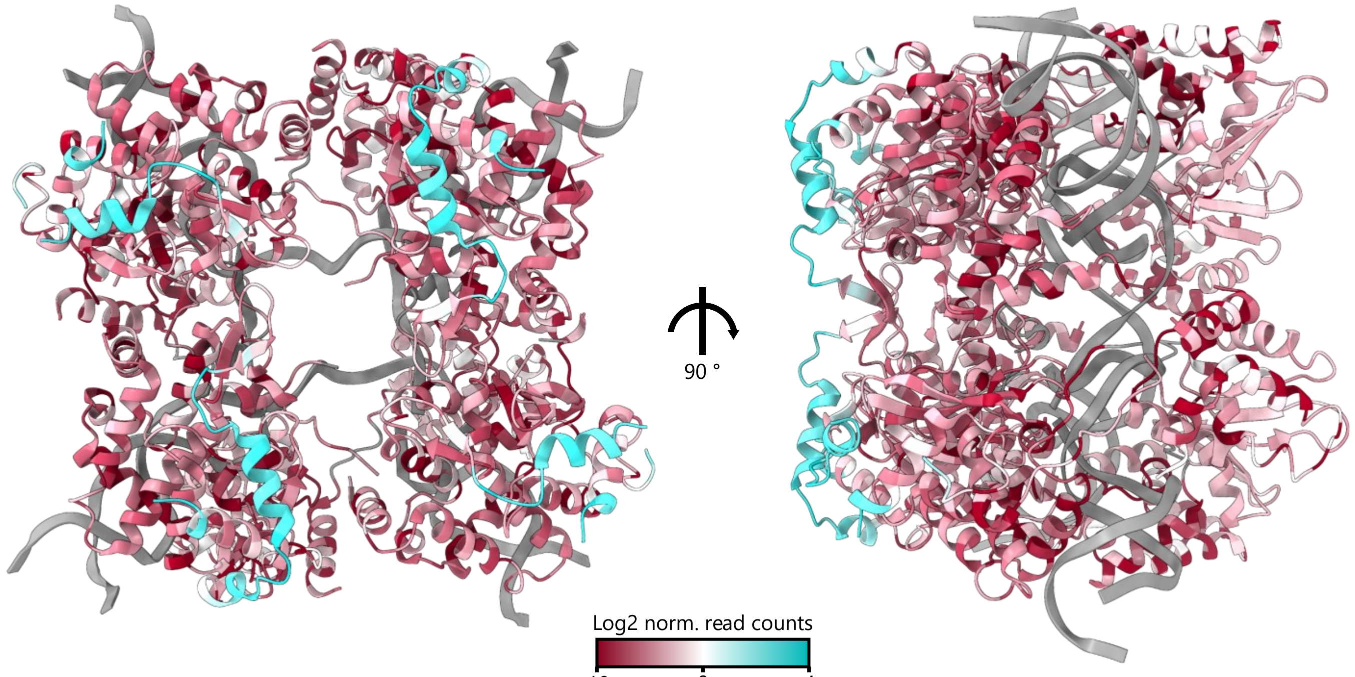


**Supplementary Fig. 4 | Enrichment scores mapped onto structures of the Flp-holliday junction complex.** PDB-ID: 1FLO.

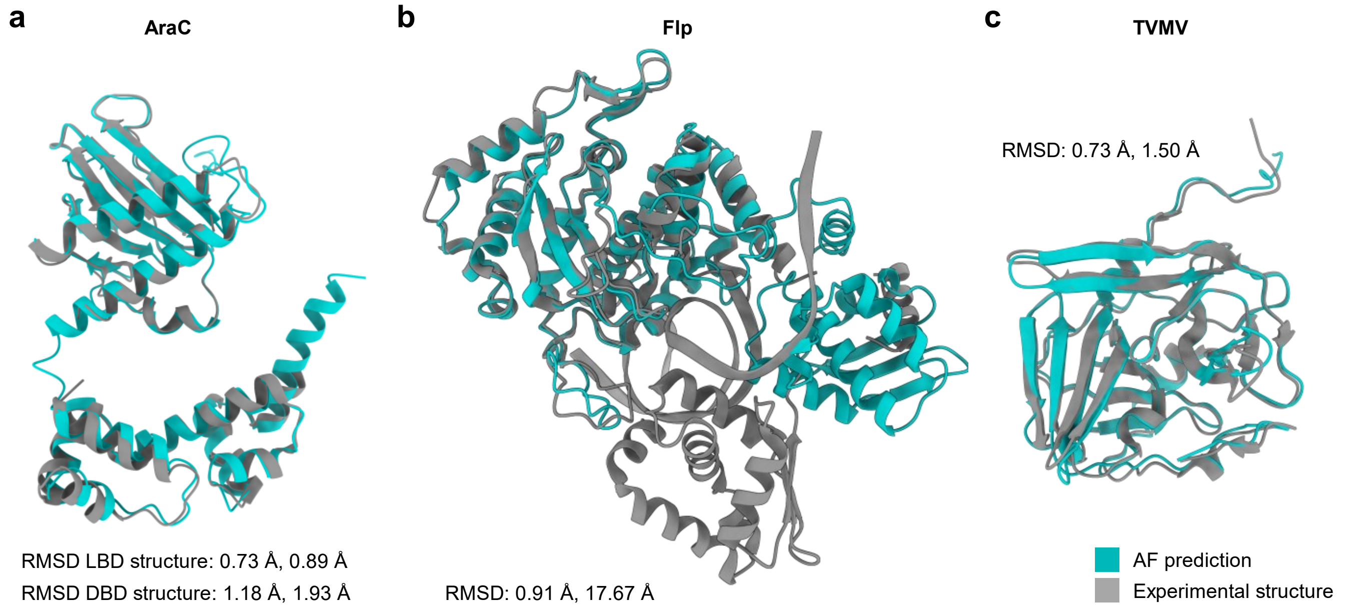


**Supplementary Fig. 5 | AlphaFold2 predictions accurately capture the structures of the candidate proteins.**

**a-c**, Structural alignments between experimentally resolved structures (grey) and AlphaFold2 predictions (green) are shown for AraC (**a**), Flp (**b**) and the TVMV protease (**c**). The RMSD of the aligned residues as well as the RMSD for all amino acids are shown. PDB-IDs: 2ARA, 2K9S, 1FLO, 3MMG.

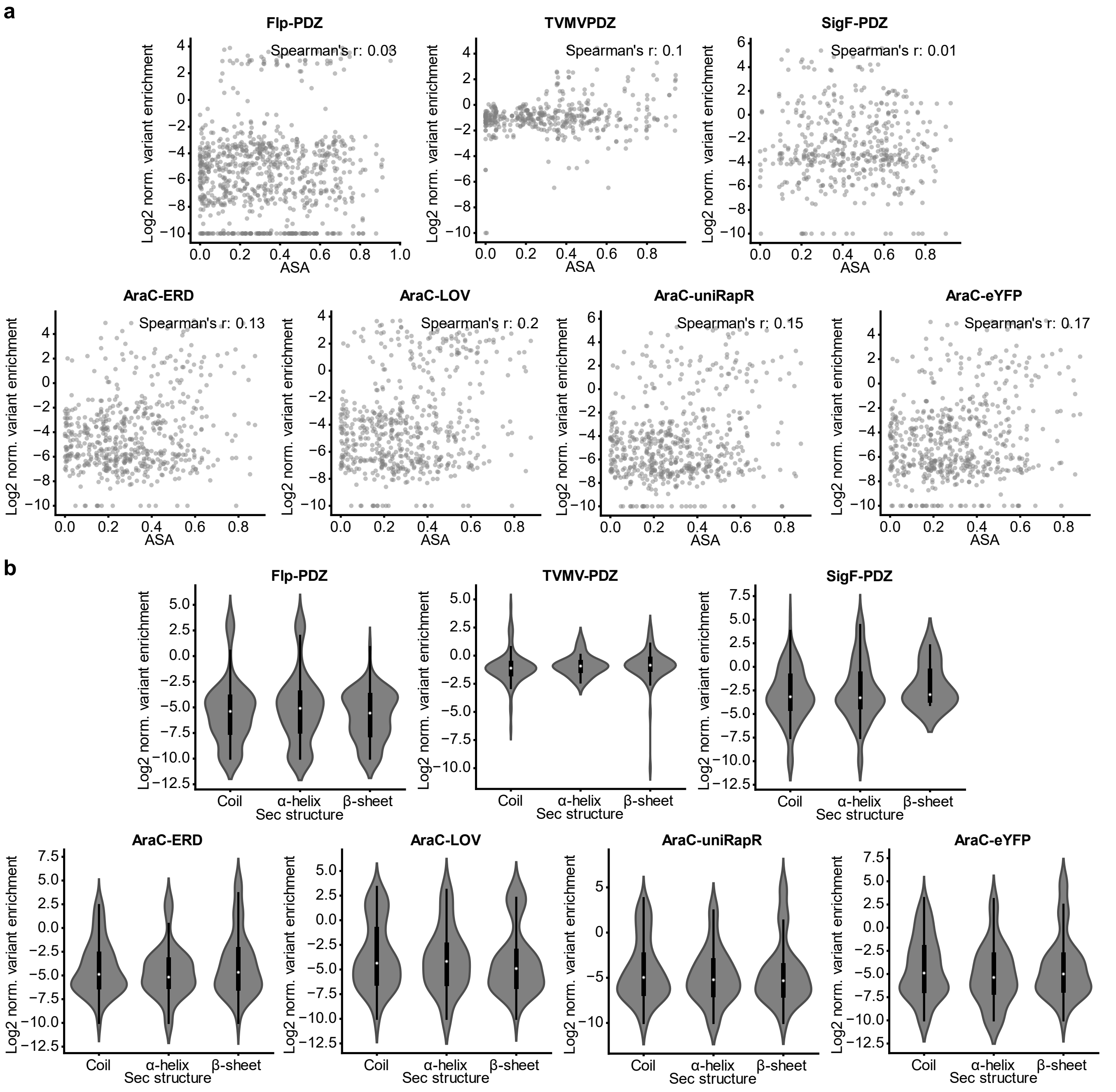


**Supplementary Fig. 6 | Correlations between the enrichment scores and surface accessibility or secondary structures.**

**a**, Scatter plot showing the relation between variant enrichment and the average surface exposed area (ASA) of the residues neighboring an insertion site. **b**, The insertion score in regions with the respective secondary structure element are shown. For each insertion site, the secondary structure assignment of the amino acid prior and after the insertion were considered. The IQR is marked by the box and the median is represented by the white dot. Whiskers extend to the 1.5-fold IQR or to the value of the smallest or largest enrichment, respectively.

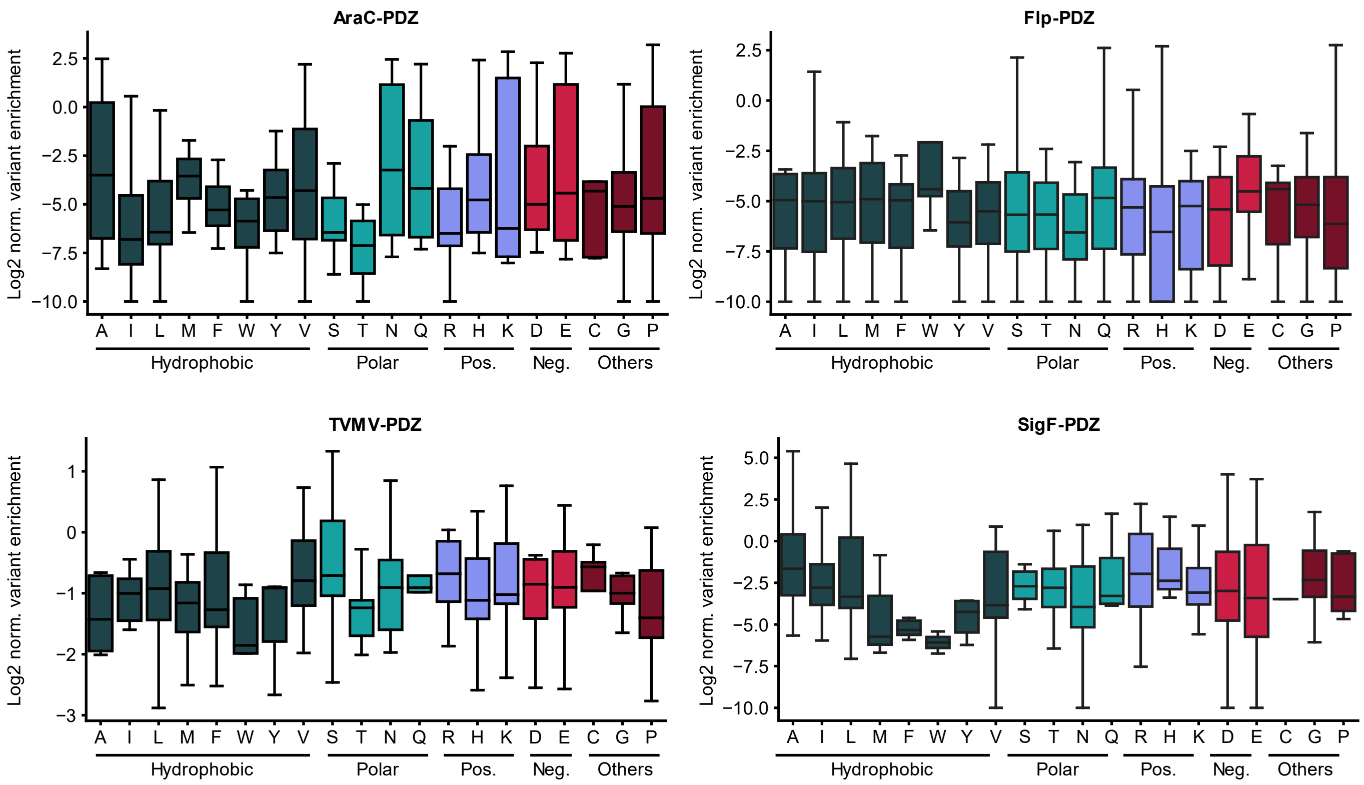


**Supplementary Fig. 7 | Successful domain insertion cannot be predicted from amino acid identity.** **a-d**, The enrichment score distribution for each amino acid is shown as boxplots for the PDZ libraries of AraC (**a**), Flp (**b**), TVMV protease (**c**) and SigF (**d**). Both residues neighboring an insertion site were taken into account for the calculations. The IQR is marked by the box and the median is represented by a line within the box. Whiskers extend to the 1.5-fold interquartile range (IQR) or to the value of the smallest or largest enrichment. Colors indicate the different amino acid categories as indicated underneath the plots. Pos., positive charged. Neg., negatively charged.

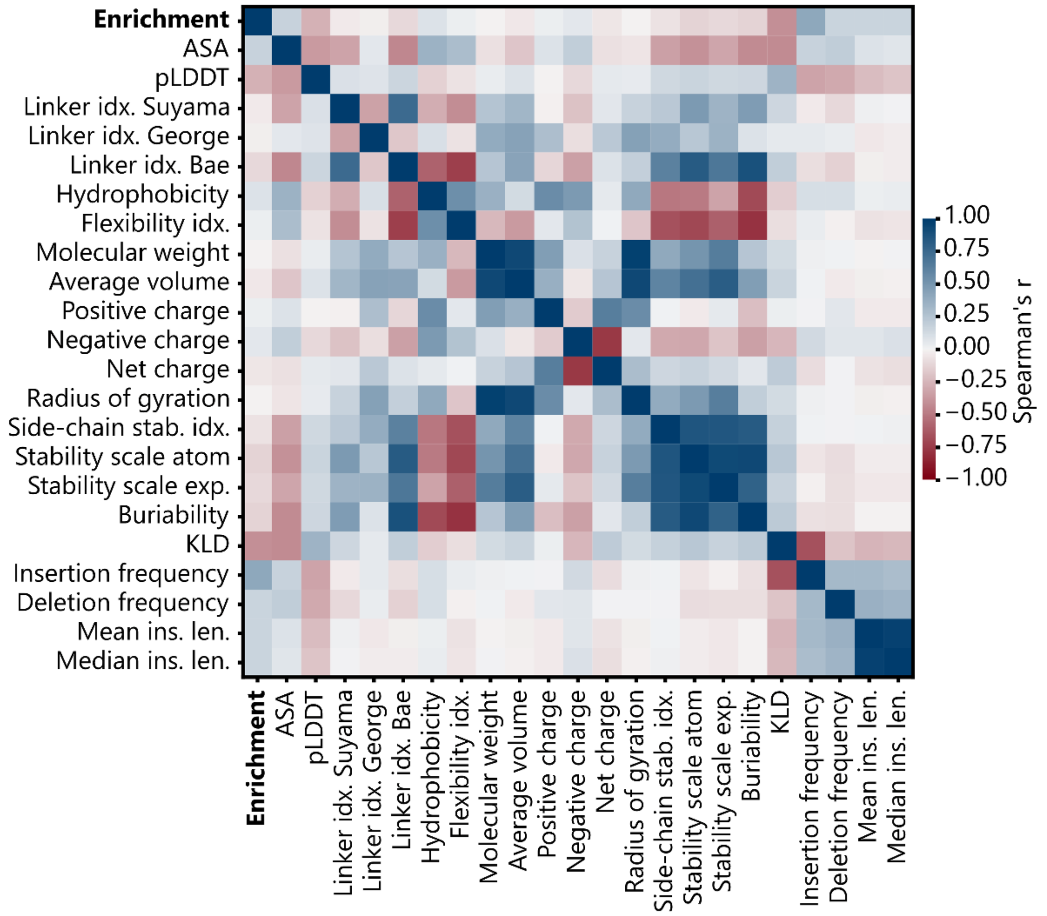


**Supplementary Fig. 8 | Heatmap of pairwise Spearman correlations between all investigated positional features.**

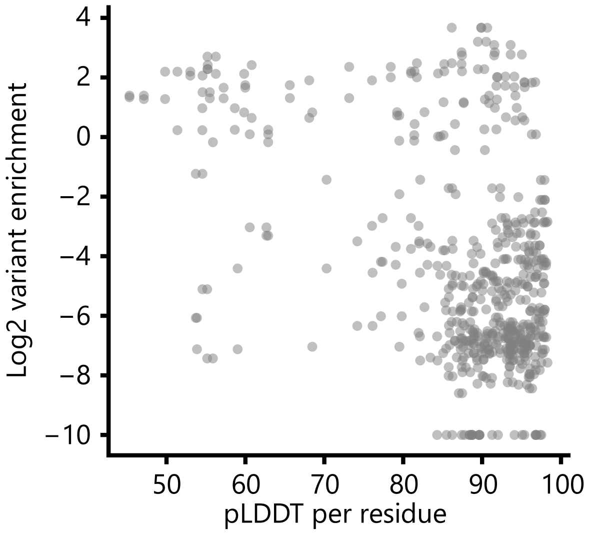


**Supplementary Fig. 9 | The position-specific pLDDT scores of wildtype AraC do not correlate with domain insertion susceptibility.**

Scatterplot of the relation between the enrichment scores of the AraC-PDZ library and the amino acid pLDDT scores from an AraC structure predicted by AF2. The corresponding Spearman's r is -0.26.

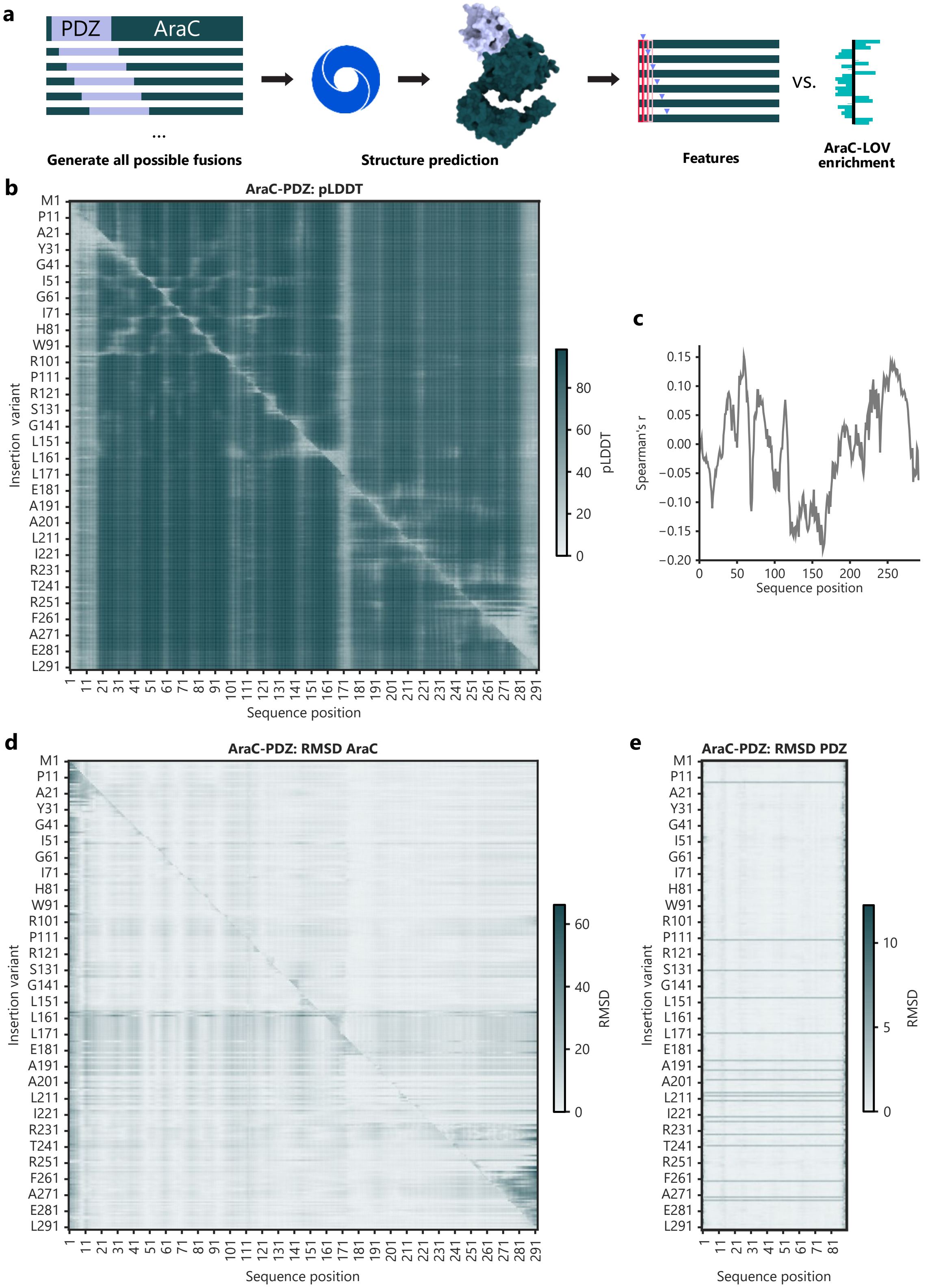


**Supplementary Fig. 10 | Correlations of AF2 structure predictions with domain insertion susceptibility.** **a**, Depiction of the structure prediction workflow. Structures for all possible insertions of the PDZ domain into AraC were generated with AF2. Structural changes at single positions in response to different insertions were then compared and correlated to the experimental enrichments. **b**, Structures of all possible PDZ insertions into AraC were predicted. The heatmap shows the pLDDT scores per position for each variant. Only AraC amino acids are depicted so that each column corresponds to pLDDT values from the same residue in different insertion variants. Rows, in turn, correspond to the different AraC-PDZ hybrids. **c**, For each amino acid position, the pLDDT scores from all variants (columns in **b**) were correlated with the corresponding enrichment scores at these positions. The resulting Spearman correlation coefficients are shown. **d-e**, The predicted AraC-PDZ structures were aligned to a predicted structure of wildtype AraC (**d**) or PDZ (**e**). The RMSDs between the wildtype and the respective part of the hybrid proteins are shown in the heatmap. Rows correspond to the different AraC-PDZ hybrids and columns to RMSD values of the same residue in different variants.

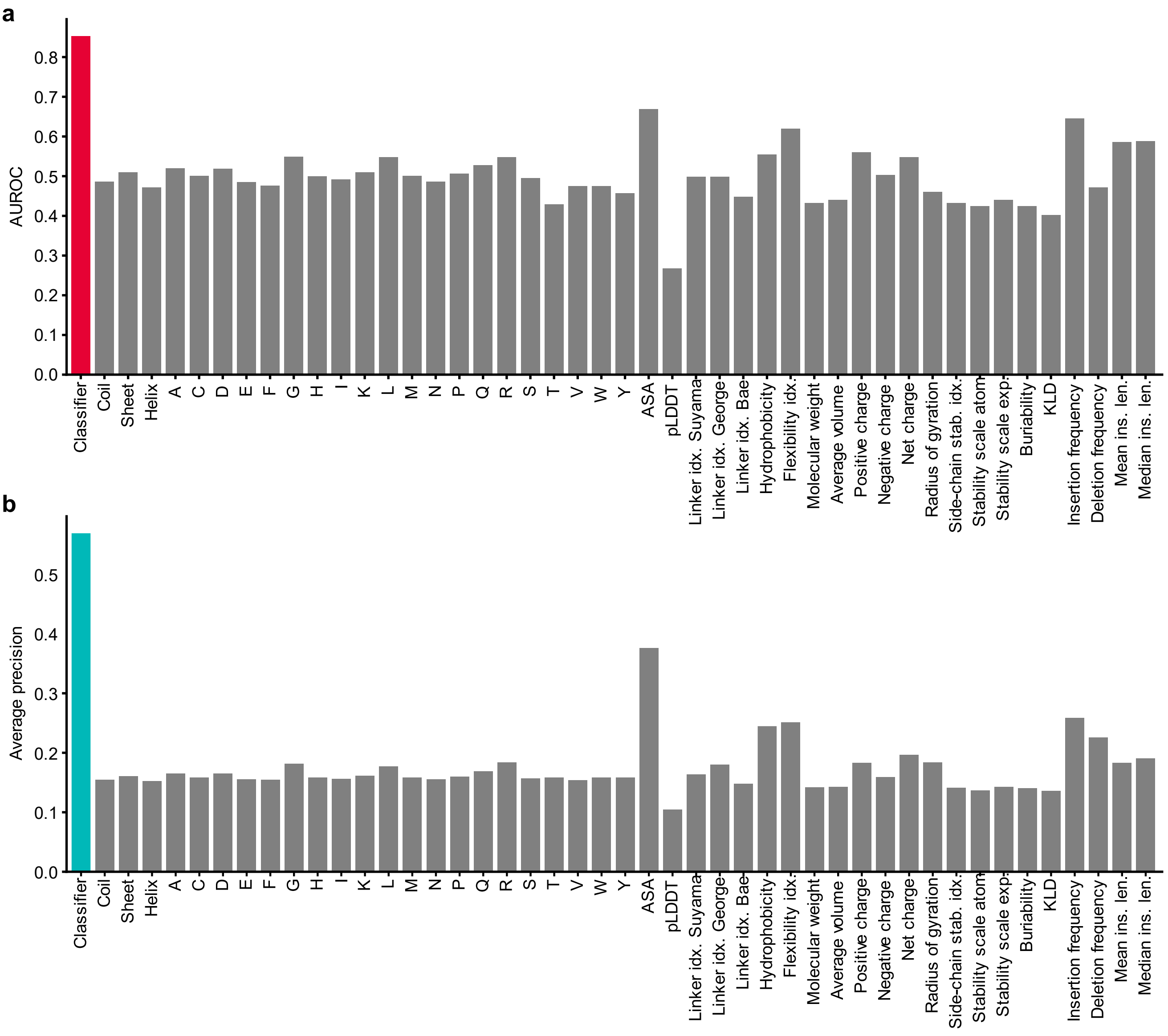


**Supplementary Fig. 11 | Full comparison of the trained classifier to baseline predictors.**

**a**, **b**, The mean AUROC (**a**) and average precision (**b**) are shown. The values were calculated on a previously withheld test set. The performance of the gradient boosting classifier is compared to all individual features.

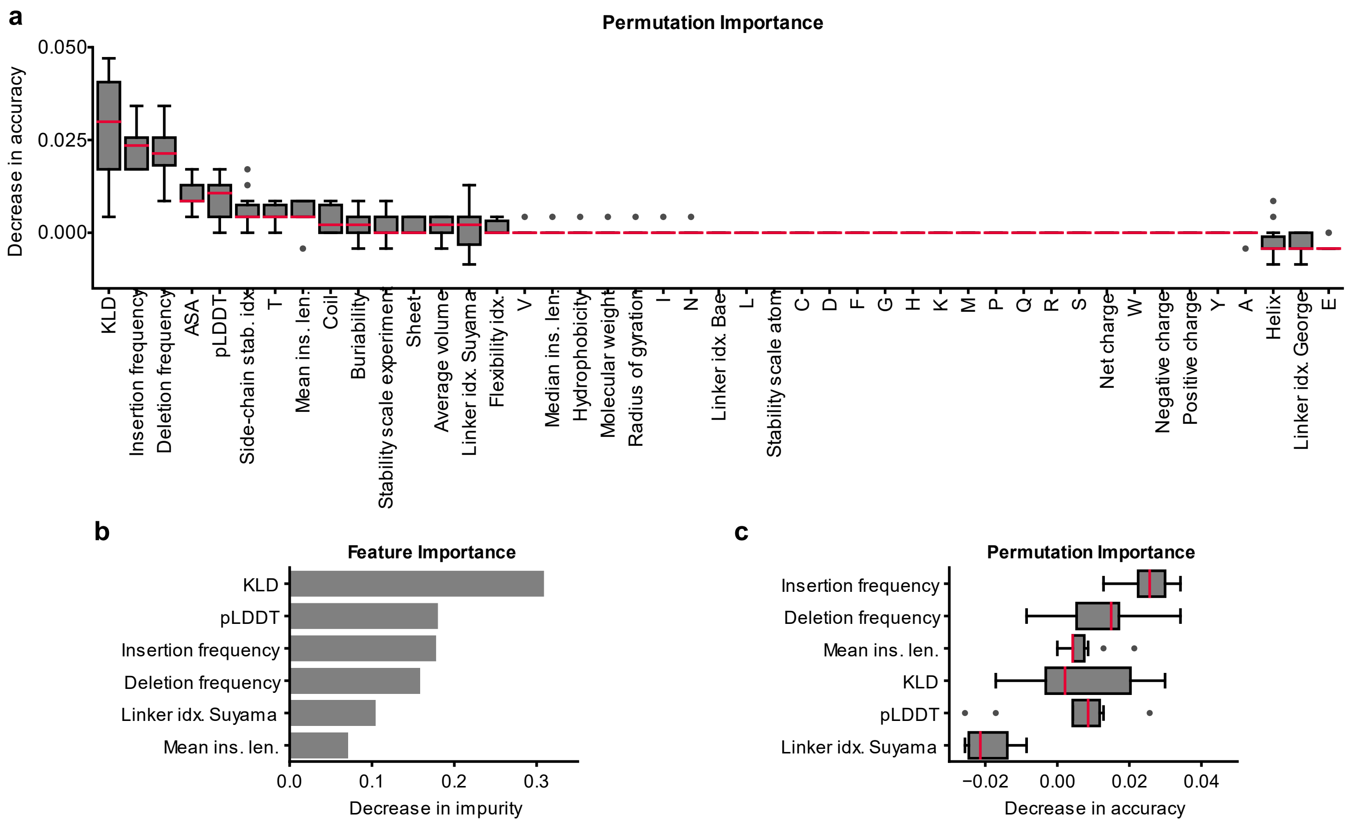


**Supplementary Fig. 12 | Alignment-derived statistics are key predictors of insertion tolerance.**

**a**, The decrease in accuracy upon random permutation of the respective features is presented for the gradient boosting model trained on the complete dataset. **b**, Bar plot indicating the Gini importance of each feature of the reduced model. **c**, The permutation importance of training features of the reduced model is shown. **a**, **c**, The results were calculated individually for each structure in the cross-validation dataset. The IQR is marked by the box and the median is represented by a red line. Whiskers extend to the 1.5-fold IQR or to the value of the smallest or largest score, respectively. Outliers are shown as points.

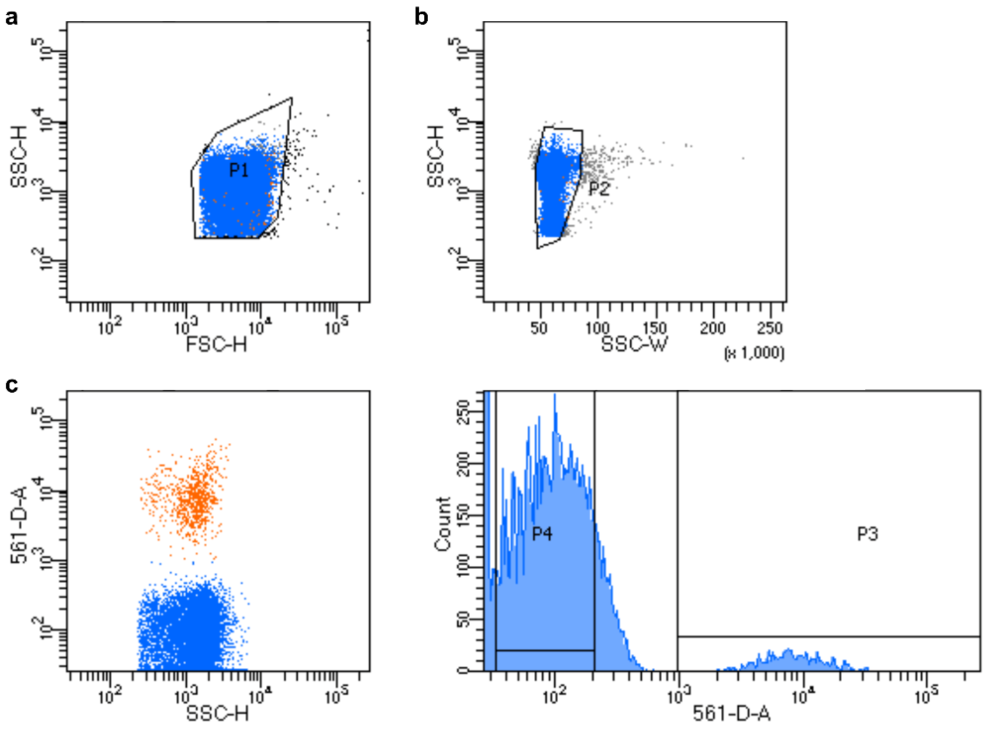


**Supplementary Fig. 13 | Gating strategy used during sorting**

**a**, Scatter plot indicating how cells were selected via their forward and side scatter. **b**, Scatter plot of side scatter height and width showing the gate that was set for the selection of singlets. **c**, The population of red fluorescent bacteria was sorted as indicated in the scatter plot and the histogram of the measured RFP fluorescence.

**Supplementary Table 1 | Constructs created and used in this study.**

| # | Name | Description. In sequential order |
|---|---|---|
| 1 | RFP reporter for AraC | BAD promoter, mRFP1, LVA degradation tag |
| 2 | RFP reporter for SigF | F1 promoter, mRFP1 |
| 3 | RFP reporter for Flp recombinase | J23102 promoter, inverted mRFP1, flanked by FRT sites |
| 4 | RFP reporter for TVMV protease (J23105 + M0051) | J23105 promoter, mRFP1, TVMV recognition site, (M0051) DAS+2 degradation tag |
| 5 | AraC | TRC promoter, AraC |
| 6 | Flp recombinase | TRC promoter, Flp recombinase |
| 7 | TVMV protease | TRC promoter, TVMV protease |
| 8 | SigF | TRC promoter, SigF |
| 9 | AraC_S170_LOV2 | TRC promoter, AraC with insertion of AsLOV2 behind S170 |
| 10 | AraC_I113_LOV2 | TRC promoter, AraC with insertion of AsLOV2 behind I113 |
| 11 | AraC_E3_PDZ | AraC with insertion of PDZ behind E3 |
| 12 | AraC_S14_PDZ | AraC with insertion of PDZ behind S14 |
| 13 | AraC_N16_PDZ | AraC with insertion of PDZ behind N16 |
| 14 | AraC_A17_PDZ | AraC with insertion of PDZ behind A17 |
| 15 | AraC_L23_PDZ | AraC with insertion of PDZ behind L23 |
| 16 | AraC_E27_PDZ | AraC with insertion of PDZ behind E27 |
| 17 | AraC_T50_PDZ | AraC with insertion of PDZ behind T50 |
| 18 | AraC_Q60_PDZ | AraC with insertion of PDZ behind Q60 |
| 19 | AraC_E63_PDZ | AraC with insertion of PDZ behind E63 |
| 20 | AraC_S112_PDZ | AraC with insertion of PDZ behind S112 |
| 21 | AraC_I113_PDZ | AraC with insertion of PDZ behind I113 |
| 22 | AraC_N116_PDZ | AraC with insertion of PDZ behind N116 |
| 23 | AraC_R121_PDZ | AraC with insertion of PDZ behind R121 |
| 24 | AraC_H129_PDZ | AraC with insertion of PDZ behind H129 |
| 25 | AraC_G143_PDZ | AraC with insertion of PDZ behind G143 |
| 26 | AraC_E165_PDZ | AraC with insertion of PDZ behind E165 |
| 27 | AraC_S170_PDZ | AraC with insertion of PDZ behind S170 |
| 28 | AraC_T241_PDZ | AraC with insertion of PDZ behind T241 |
| 29 | AraC_D286_PDZ | AraC with insertion of PDZ behind D286 |
| 30 | AraC_E3_LOV2 | AraC with insertion of LOV2 behind E3 |
| 31 | AraC_S14_LOV2 | AraC with insertion of LOV2 behind S14 |
| 32 | AraC_N16_LOV2 | AraC with insertion of LOV2 behind N16 |
| 33 | AraC_A17_LOV2 | AraC with insertion of LOV2 behind A17 |
| 34 | AraC_L23_LOV2 | AraC with insertion of LOV2 behind L23 |
| 35 | AraC_E27_LOV2 | AraC with insertion of LOV2 behind E27 |
| 36 | AraC_T50_LOV2 | AraC with insertion of LOV2 behind T50 |
| 37 | AraC_Q60_LOV2 | AraC with insertion of LOV2 behind Q60 |
| 38 | AraC_E63_LOV2 | AraC with insertion of LOV2 behind E63 |
| 39 | AraC_S112_LOV2 | AraC with insertion of LOV2 behind S112 |
| 40 | AraC_I113_LOV2 | AraC with insertion of LOV2 behind I113 |
| 41 | AraC_N116_LOV2 | AraC with insertion of LOV2 behind N116 |
| 42 | AraC_R121_LOV2 | AraC with insertion of LOV2 behind R121 |
| 43 | AraC_H129_LOV2 | AraC with insertion of LOV2 behind H129 |
| 44 | AraC_G143_LOV2 | AraC with insertion of LOV2 behind G143 |

| | | |
|---|---|---|
| 45 | AraC_E165_LOV2 | AraC with insertion of LOV2 behind E165 |
| 46 | AraC_S170_LOV2 | AraC with insertion of LOV2 behind S170 |
| 47 | AraC_T241_LOV2 | AraC with insertion of LOV2 behind T241 |
| 48 | AraC_D286_LOV2 | AraC with insertion of LOV2 behind D286 |
| 49 | AraC_E3_eYFP | AraC with insertion of eYFP behind E3 |
| 50 | AraC_N16_eYFP | AraC with insertion of eYFP behind N16 |
| 51 | AraC_L23_eYFP | AraC with insertion of eYFP behind L23 |
| 52 | AraC_T50_eYFP | AraC with insertion of eYFP behind T50 |
| 53 | AraC_Q60_eYFP | AraC with insertion of eYFP behind Q60 |
| 54 | AraC_I113_eYFP | AraC with insertion of eYFP behind I113 |
| 55 | AraC_N116_eYFP | AraC with insertion of eYFP behind N116 |
| 56 | AraC_E165_eYFP | AraC with insertion of eYFP behind E165 |
| 57 | AraC_S170_eYFP | AraC with insertion of eYFP behind S170 |
| 58 | AraC_T241_eYFP | AraC with insertion of eYFP behind T241 |
| 59 | AraC_E3_ERD | AraC with insertion of ERD behind E3 |
| 60 | AraC_N16_ERD | AraC with insertion of ERD behind N16 |
| 61 | AraC_L23_ERD | AraC with insertion of ERD behind L23 |
| 62 | AraC_T50_ERD | AraC with insertion of ERD behind T50 |
| 63 | AraC_Q60_ERD | AraC with insertion of ERD behind Q60 |
| 64 | AraC_I113_ERD | AraC with insertion of ERD behind I113 |
| 65 | AraC_N116_ERD | AraC with insertion of ERD behind N116 |
| 66 | AraC_E165_ERD | AraC with insertion of ERD behind E165 |
| 67 | AraC_S170_ERD | AraC with insertion of ERD behind S170 |
| 68 | AraC_T241_ERD | AraC with insertion of ERD behind T241 |
| 69 | AraC_E3_uniRapR | AraC with insertion of uniRapR behind E3 |
| 70 | AraC_N16_uniRapR | AraC with insertion of uniRapR behind N16 |
| 71 | AraC_L23_uniRapR | AraC with insertion of uniRapR behind L23 |
| 72 | AraC_T50_uniRapR | AraC with insertion of uniRapR behind T50 |
| 73 | AraC_Q60_uniRapR | AraC with insertion of uniRapR behind Q60 |
| 74 | AraC_I113_uniRapR | AraC with insertion of uniRapR behind I113 |
| 75 | AraC_N116_uniRapR | AraC with insertion of uniRapR behind N116 |
| 76 | AraC_E165_uniRapR | AraC with insertion of uniRapR behind E165 |
| 77 | AraC_S170_uniRapR | AraC with insertion of uniRapR behind S170 |
| 78 | AraC_T241_uniRapR | AraC with insertion of uniRapR behind T241 |
| 79 | TVMV_L5_PDZ | TVMV with insertion of PDZ behind L5 |
| 80 | TVMV_D11_PDZ | TVMV with insertion of PDZ behind D11 |
| 81 | TVMV_G37_PDZ | TVMV with insertion of PDZ behind G37 |
| 82 | TVMV_I42_PDZ | TVMV with insertion of PDZ behind I42 |
| 83 | TVMV_L56_PDZ | TVMV with insertion of PDZ behind L56 |
| 84 | TVMV_T105_PDZ | TVMV with insertion of PDZ behind T105 |
| 85 | TVMV_S121_PDZ | TVMV with insertion of PDZ behind S121 |
| 86 | TVMV_H143_PDZ | TVMV with insertion of PDZ behind H143 |
| 87 | TVMV_F187_PDZ | TVMV with insertion of PDZ behind F187 |
| 88 | TVMV_D193_PDZ | TVMV with insertion of PDZ behind D193 |
| 89 | TVMV_W198_PDZ | TVMV with insertion of PDZ behind W198 |
| 90 | TVMV_F204_PDZ | TVMV with insertion of PDZ behind F204 |
| 91 | TVMV_I209_PDZ | TVMV with insertion of PDZ behind I209 |
| 92 | Flp_L15_PDZ | Flp with insertion of PDZ behind L15 |

| | | |
|---|---|---|
| 93 | Flp_C42_PDZ | Flp with insertion of PDZ behind C42 |
| 94 | Flp_D115_PDZ | Flp with insertion of PDZ behind D115 |
| 95 | Flp_S129_PDZ | Flp with insertion of PDZ behind S129 |
| 96 | Flp_L151_PDZ | Flp with insertion of PDZ behind L151 |
| 97 | Flp_I239_PDZ | Flp with insertion of PDZ behind I239 |
| 98 | Flp_N290_PDZ | Flp with insertion of PDZ behind N290 |
| 99 | Flp_W330_PDZ | Flp with insertion of PDZ behind W330 |
| 100 | Flp_S397_PDZ | Flp with insertion of PDZ behind S397 |
| 101 | Flp_Y403_PDZ | Flp with insertion of PDZ behind Y403 |
| 102 | AraC_E3I | AraC with E3I mutation |
| 103 | AraC_A28Y | AraC with A28Y mutation |
| 104 | AraC_T50S | AraC with T50S mutation |
| 105 | AraC_I113D | AraC with I113D mutation |
| 106 | AraC_G141D | AraC with G141D mutation |
| 107 | AraC_G141V | AraC with G141V mutation |
| 108 | AraC_G141Y | AraC with G141Y mutation |
| 109 | AraC_E165I | AraC with E165I mutation |
| 110 | AraC_T241C | AraC with T241C mutation |
| 111 | AraC_V284F | AraC with V284F mutation |
| 112 | AraC_V284I | AraC with V284I mutation |
| 113 | AraC_I113_LOV_E3I | AraC_I113_LOV with E3I mutation |
| 114 | AraC_I113_LOV_A28Y | AraC_I113_LOV with A28Y mutation |
| 115 | AraC_I113_LOV_T50S | AraC_I113_LOV with T50S mutation |
| 116 | AraC_I113_LOV_G141D | AraC_I113_LOV with G141D mutation |
| 117 | AraC_I113_LOV_G141V | AraC_I113_LOV with G141V mutation |
| 118 | AraC_I113_LOV_G141Y | AraC_I113_LOV with G141Y mutation |
| 119 | AraC_I113_LOV_E165I | AraC_I113_LOV with E165I mutation |
| 120 | AraC_I113_LOV_T241C | AraC_I113_LOV with T241C mutation |
| 121 | AraC_I113_LOV_V284F | AraC_I113_LOV with V284F mutation |
| 122 | AraC_I113_LOV_V284I | AraC_I113_LOV with V284I mutation |
| 123 | AraC_S170_LOV_E3I | AraC_S170_LOV with E3I mutation |
| 124 | AraC_S170_LOV_A28Y | AraC_S170_LOV with A28Y mutation |
| 125 | AraC_S170_LOV_T50S | AraC_S170_LOV with T50S mutation |
| 126 | AraC_S170_LOV_I113D | AraC_S170_LOV with I113D mutation |
| 127 | AraC_S170_LOV_G141D | AraC_S170_LOV with G141D mutation |
| 128 | AraC_S170_LOV_G141V | AraC_S170_LOV with G141V mutation |
| 129 | AraC_S170_LOV_G141Y | AraC_S170_LOV with G141Y mutation |
| 130 | AraC_S170_LOV_T241C | AraC_S170_LOV with T241C mutation |
| 131 | AraC_S170_LOV_V284F | AraC_S170_LOV with V284F mutation |
| 132 | AraC_S170_LOV_V284I | AraC_S170_LOV with V284I mutation |

**Supplementary Table 2 | Amino acid sequences of the used proteins and insert domains.** Tags linked to the proteins are marked in bold.

| Protein | Sequence |
|---|---|
| AraC | MSAEAQNDPLLPGYSFNAHLVAGLTPIEANGYLDFFIDRPLGMKGYILNLTIRGQGVVKNQGR<br>EFVCRPGDILLFPPGEIHHYGRHPEAREWYHQWVYFRPRAYWHEWLNWPSIFANTGFFRPDEA<br>HQPHFSDLFGQIINAGQGEGRYSELLAINLLEQLLLRRMEAINESLHPPMDNRVREACQYISD<br>HLADSNFDIASVAQHVCLSPSRLSHLFRQQLGISVLSWREDQRISQAKLLLSTTRMPIATVGR<br>NVGFDDQLYFSRVFKKCTGASPSEFRAGCEEKVNDVAVKLS**GHHHHHH** |
| *As*LOV2 | LATTLERIEKNFVITDPRLPDNPIIFASDSFLQLTEYSREEILGRNCRFLQGPETDRATVRKI<br>RDAIDNQTEVTVQLINYTKSGKKFWNLFHLQPMRDQKGDVQYFIGVQLDGTEHVRDAAEREGV<br>MLIKKTAENIDEAAK |
| ER domain | GPLDNSLALSLTADQMVSALLDAEPPILYSEYDPTRPFSEASMMGLLTNLADRELVHMINWAK<br>RVPGFVDLTLHDQVHLLECAWLEILMIGLVWRSMEHPGKLLFAPNLLLDRNQGKCVEGMVEIF<br>DMLLATSSRFRMMNLQGEEFVCLKSIILLNSGVYTFLSSTLKSLEEKDHIHRVLDKITDTLIH<br>LMAKAGLTLQQQHQRLAQLLLILSHIRHMSNKGMEHLYSMKCKNVVPLYDLLLEMLDAHRLHA<br>PGSEL |
| eYFP | VSKGEELFTGVVPILVELDGDVNGHKFSVSGEGEGDATYGKLTLKFICTTGKLPVPWPTLVTT<br>FGYGLQCFARYPDHMKQHDFFKSAMPEGYVQERTIFFKDDGNYKTRAEVKFEGDTLVNRIELK<br>GIDFKEDGNILGHKLEYNYNSHNVYIMADKQKNGIKVNFKIRHNIEDGSVQLADHYQQNTPIG<br>DGPVLLPDNHYLSYQSKLSKDPNEKRDHMVLLEFVTAAGITLGMDELYK |
| Flp recombinase | MSPQFGILCKTPPKVLVRQFVERFERPSGEKIALCAAELTYLCWMITHNGTAIKRATFMSYNT<br>IISNSLSFDIVNKSLQFKYKTQKATILEASLKKLIPAWEFTIIPYYGQKHQSDITDIVSSLQL<br>QFESSEEADKGNSHSKKMLKALLSEGESIWEITEKILNSFEYTSRFTKTKTLYQFLFLATFIN<br>CGRFSDIKNVDPKSFKLVQNKYLGVIIQCLVTETKTSVSRHIYFFSARGRIDPLVYLDEFLRN<br>SEPVLKRVNRTGNSSSNKQEYQLLKDNLVRSYNKALKKNAPYSIFAIKNGPKSHIGRHLMTSF<br>LSMKGLTELTNVVGNWSDKRASAVARTTYTHQITAIPDHYFALVSRYYAYDPISKEMIALKDE<br>TNPIEEWQHIEQLKGSAEGSIRYPAWNGIISQEVLDYLSSYINRRI**SGHHHHHH** |
| mRFP1 | MASSEDVIKEFMRFKVRMEGSVNGHEFEIEGEGEGRPYEGTQTAKLKVTKGGPLPFAWDILSP<br>QFQYGSKAYVKHPADIPDYLKLSFPEGFKWERVMNFEDGGVVTVTQDSSLQDGEFIYKVKLRG<br>TNFPSDGPVMQKKTMGWEASTERMYPEDGALKGEIKMRLKLKDGGHYDAEVKTTYMAKKPVQL<br>PGAYKTDIKLDITSHNEDYTIVEQYERAEGRHSTGA |
| PDZ domain | RRRVTVRKADAGGLGISIKGGRENKMPILISKIFKGLAADQTEALFVGDAILSVNGEDLSSAT<br>HDEAVQALKKTGKEVVLEVKYMK |
| SigF | MSDVEVKKNGKNAQLKDHEVKELIKQSQNGDQQARDLLIEKNMRLVWSVVQRFLNRGYEPDDL<br>FQIGCIGLLKSVDKFDLTYDVRFSTYAVPMIIGEIQRFIRDDGTVKVSRSLKELGNKIRRAKD<br>ELSKTLGRVPTVQEIADHLEIEAEDVVLAQEAVRAPSSIHETVYENDGDPITLLDQIADNSEE<br>KWFDKIALKEAISDLEEREKLIVYLRYYKDQTQSEVAERLGISQVQVSRLEKKILKQIKVQMD<br>HTDG |
| TVMV protease | MSSKALLKGVRDFNPISACVCLLENSSDGHSERLFGIGFGPYIIANQHLFRRNNGELTIKTMH<br>GEFKVKNSTQLQMKPVEGRDIIVIKMAKDFPPFPQKLKFRQPTIKDRVCMVSTNFQQKSVSSL<br>VSESSHIVHKEDTSFWQHWITTKDGQCGSPLVSIIDGNILGIHSLTHTTNGSNYFVEFPEKFV<br>ATYLDAADGWCKNWKFNADKISWGSFTLVE |
| uniRapR | TCVVHYTGMLEDGKKFDSSRDRNKPFKFMLGKQEVIRGWEEGVAQMSVGQRAKLTISPDYAYG<br>ATGHGSGSGSGVKDLLQAWDLYYHVFRRISGPPGPGSGLWHEMWHEGLEEASRLYFGERNVKG<br>MFEVLEPLHAMMERGPQTLKETSFNQAYGRDLMEAQEWCRKYMKSGSSGGSGSGIIPPHATLV<br>FDVELLKLE |

**Supplementary Table 3 | PDB IDs of protein structures shown and used in this study.**

| Structure | PDB-ID | Reference |
|---|---|---|
| AraC, apo-form | 2ARA | [14] |
| AraC, complexed with L-arabinose | 2ARC | [14] |
| AraC, DBD | 2K9S | [15] |
| *As*LOV2 domain | 2V0W, 2V0U | [16] |
| ERD | 1A52 | [17] |
| eYFP, F165G | 6ZQO | [18] |
| Flp recombinase | 1FLO | [19] |
| PDZ | 1Z86 | [20] |
| Rob transcription factor | 1D5Y | [21] |
| TVMV protease | 3MMG | [22] |
| uniRapR | 1FAP | [23] |

### Supplementary References


1. Akdel, M. *et al.* A structural biology community assessment of AlphaFold2 applications. *Nat Struct Mol Biol* 29, 1056–1067 (2022).
2. Saldaño, T. *et al.* Impact of protein conformational diversity on AlphaFold predictions. *Bioinformatics* 38, 2742–2748 (2022).
3. Wilson, C. J., Choy, W.-Y. & Karttunen, M. AlphaFold2: A Role for Disordered Protein/Region Prediction? *Int J Mol Sci* 23, 4591 (2022).
4. Jumper, J. *et al.* Highly accurate protein structure prediction with AlphaFold. *Nature* 596, 583–589 (2021).
5. Nielsen, A. A. K. *et al.* Genetic circuit design automation. *Science* 352, 7341, (2016).
6. Green, A. A. *et al.* Complex cellular logic computation using ribocomputing devices. *Nature* 548, 117–121 (2017).
7. Chen, Z. *et al.* De novo design of protein logic gates. *Science* 368, 78–84 (2020).
8. Gao, X. J., Chong, L. S., Kim, M. S. & Elowitz, M. B. Programmable protein circuits in living cells. *Science* 361, 1252–1258 (2018).
9. Chen, Z., Linton, J. M., Zhu, R. & Elowitz, M. B. A synthetic protein-level neural network in mammalian cells. Preprint at https://doi.org/10.1101/2022.07.10.499405 (2022).
10. Vishweshwaraiah, Y. L., Chen, J., Chirasani, V. R., Tabdanov, E. D. & Dokholyan, N. V. Two-input protein logic gate for computation in living cells. *Nat Commun* 12, 6615 (2021).
11. Dietler, J. *et al.* A Light-Oxygen-Voltage Receptor Integrates Light and Temperature. *Journal of Molecular Biology* 433, 167107 (2021).
12. Ceroni, F. *et al.* Burden-driven feedback control of gene expression. *Nat Methods* 15, 387–393 (2018).
13. Eldar, A. & Elowitz, M. B. Functional roles for noise in genetic circuits. *Nature* 467, 167–173 (2010).
14. Soisson, S. M., MacDougall-Shackleton, B., Schleif, R. & Wolberger, C. Structural Basis for Ligand-Regulated Oligomerization of AraC. *Science* 276, 421–425 (1997).
15. Rodgers, M. E. & Schleif, R. Solution structure of the DNA binding domain of AraC protein. *Proteins: Structure, Function, and Bioinformatics* 77, 202–208 (2009).
16. Halavaty, A. S. & Moffat, K. N- and C-terminal flanking regions modulate light-induced signal transduction in the LOV2 domain of the blue light sensor phototropin 1 from Avena sativa. *Biochemistry* 46, 14001–14009 (2007).

17. Tanenbaum, D. M., Wang, Y., Williams, S. P. & Sigler, P. B. Crystallographic comparison of the estrogen and progesterone receptor's ligand binding domains. *Proceedings of the National Academy of Sciences of the United States of America* 95, 5998–6003 (1998).
18. Pletneva, N. V. *et al.* Amino acid residue at the 165th position tunes EYFP chromophore maturation. A structure-based design. *Computational and Structural Biotechnology Journal* 19, 2950–2959 (2021).
19. Chen, Y., Narendra, U., Iype, L. E., Cox, M. M. & Rice, P. A. Crystal structure of a Flp recombinase-Holliday junction complex: assembly of an active oligomer by helix swapping. *Mol Cell* 6, 885–897 (2000).
20. Yan, J. *et al.* Structure of the split PH domain and distinct lipid-binding properties of the PH–PDZ supramodule of α-syntrophin. *The EMBO Journal* 24, 3985–3995 (2005).
21. Kwon, H. J., Bennik, M. H. J., Demple, B. & Ellenberger, T. Crystal structure of the Escherichia coli Rob transcription factor in complex with DNA. *Nat Struct Mol Biol* 7, 424–430 (2000).
22. Sun, P., Austin, B. P., Tözsér, J. & Waugh, D. S. Structural determinants of tobacco vein mottling virus protease substrate specificity: Structure of TVMV Protease/Substrate Complex. *Protein Science* 19, 2240–2251 (2010).
23. Choi, J., Chen, J., Schreiber, S. L. & Clardy, J. Structure of the FKBP12-Rapamycin Complex Interacting with Binding Domain of Human FRAP. *Science* 273, 239–242 (1996).
